# ENGRAILED-1 transcription factor has a paracrine neurotrophic activity on adult spinal α-motoneurons

**DOI:** 10.1101/2022.08.16.504081

**Authors:** Mélanie Lebœuf, Stephanie E. Vargas-Abonce, Eugénie Pezé-Hedsieck, Edmond Dupont, Lucia Jimenez-Alonso, Kenneth L. Moya, Alain Prochiantz

**Affiliations:** Center for Interdisciplinary Research in Biology, Collège de France, CNRS, INSERM, Université PSL, Paris, France; BrainEver SAS, Paris France

**Keywords:** ALS, ENGRAILED1, Motoneurons, Neurodegeneration, Paracrine

## Abstract

Several homeoprotein transcription factors transfer between cells and regulate gene expression, protein translation, and chromatin organization in recipient cells. ENGRAILED-1 is one such homeoprotein expressed in spinal V1 interneurons synapsing on α-motoneurons. Neutralizing extracellular ENGRAILED-1 by expressing a secreted single-chain antibody blocks its capture by spinal motoneurons resulting in α-motoneurons loss and limb weakness. A similar but stronger phenotype is observed in the *Engrailed-1* heterozygote mouse, confirming that ENGRAILED-1 exerts a paracrine neurotrophic activity on spinal cord α-motoneurons. Intrathecal injection of ENGRAILED-1 leads to its specific internalization by spinal motoneurons and has long-lasting protective effects against neurodegeneration and weakness. Midbrain dopaminergic neurons express *Engrailed-1* and, similarly to spinal cord α-motoneurons, degenerate in the heterozygote. By identifying genes expressed in spinal cord motoneurons also showing modified expression in mouse *Engrailed-1* heterozygote midbrain neurons, we identified p62/SQTSM1 as an age marker in spinal cord motoneurons with increased expression during aging, in the *Engrailed-1* heterozygote and upon extracellular ENGRAILED-1 neutralization. We conclude that ENGRAILED-1 is a regulator of motoneuron ageing with non-cell autonomous neurotrophic activity.

## INTRODUCTION

Homeoprotein (HP) transcription factors are key transcriptional regulators with well-established developmental functions, including control of morphogenesis, lineage decisions, and cell differentiation (Gehring, 1987; Holland & Takahashi, 2005). In addition to their classical cell autonomous activities, non-cell autonomous developmental functions based on HP intercellular transfer were identified for ENGRAILED-1 (EN1), EN2, VAX1, OTX2, and PAX6. Early in development, paracrine ENGRAILED induces anterior cross vein formation in the fly wing disk and shapes the zebrafish optic tectum (Layalle *et al*, 2011; Rampon *et al*, 2015; Amblard *et al*, 2020b). ENGRAILED-2 and VAX1 respectively regulate retinal ganglion cell (RGC) axon guidance in the tectum and decussation at the optic chiasma (Wizenmann *et al*, 2009; Brunet *et al*, 2005; Yoon *et al*, 2012; Kim *et al*, 2014), while PAX6 regulates the size of the zebrafish eye anlagen, enhances oligodendrocyte progenitor cell migration in the chick neural tube, and guides Cajal-Retzius cell migration in the mouse neuroepithelium (Di Lullo *et al*, 2011; Kaddour *et al*, 2019; Lesaffre *et al*, 2007). At late developmental stages and in the adult, OTX2 non-cell autonomous activity regulates cerebral cortex plasticity (Sugiyama *et al*, 2008; Di Nardo *et al*, 2020), inner retinal physiological functions and RGC survival (Torero-Ibad *et al*, 2020; Torero-Ibad *et al*, 2011; Pensieri *et al*, 2021). This novel signaling pathway, involving unconventional secretion and internalization with direct access to the cytoplasm, is likely common to a larger number of HPs since the transfer domains are highly conserved between most HPs (Prochiantz & Joliot, 2003). The finding that more than 150 HPs can transfer between cells, *in vitro* and in the embryonic brain, even though without identified functions for most of them, lends weight to this hypothesis (Lee *et al*, 2019).

Several vertebrate HPs remain expressed throughout life, but their adult physiological functions are still poorly understood. In the context of neurodegenerative diseases, EN1 is of particular interest since it is expressed in the mesencephalic dopaminergic (mDA) neurons of the Ventral Tegmental Area (VTA) and Substantia Nigra pars compacta (SNpc) (Sgadò *et al*, 2006; Di Nardo *et al*, 2007), two cell populations that degenerate in Parkinson’s Disease (PD), and promotes their survival in PD animal models (Di Nardo *et al*, 2018). In the *En1^+/-^* mouse (*En1-*Het), mDA neurons from the VTA and SNpc undergo progressive retrograde degeneration leading to a loss of 20% and 40% of their initial number, respectively (Sgadò *et al*, 2006; Sonnier *et al*, 2007; Nordström *et al*, 2015). Conversely, EN1 injection in the midbrain and its subsequent internalization by mDA neurons, thanks to its transduction properties, prevents their degeneration in rodent and macaque PD models (Alvarez-Fischer *et al*, 2011; Thomasson *et al*, 2019). Since mDA neurons express EN1, the latter observations reflect a cell-autonomous pro-survival activity of EN1. This incited us to search for other neurons that may require EN1 for their survival, either expressing *En1* or in the vicinity of *En1*-expressing cells and thus potentially exposed to secreted EN1. In the spinal cord, inhibitory V1 interneurons, including Renshaw cells (RCs), have been shown to express *En1*, and α-motoneurons (αMNs) are candidate non-cell autonomous targets owing to their direct contact with *En1*-expressing V1 interneurons (Wenner *et al*, 2000; Alvarez *et al*, 2005; Higashijima *et al*, 2004).

We confirm here that *En1* is expressed in adult spinal cord V1 interneurons and now report that these interneurons do not degenerate in the *En1*-Het mouse. In contrast, αMNs of *En1*-Het mice show retrograde degeneration associated with progressive limb strength loss. This phenotype is in part the consequence of local EN1 non-cell autonomous activity since it can be partially mimicked in wild-type (WT) mice following EN1 neutralization in the spinal cord extracellular space. Furthermore, EN1 intrathecally injected at 3 months gains direct access to MNs and, in *En1*-Het mice, a single injection restores limb strength and prevents αMN death for at least 2.5 months. This demonstrates a novel non-cell autonomous function of EN1 in the adult CNS and suggests that this HP may be of interest for developing novel therapeutic strategies and targets for diseases involving αMN degeneration.

## RESULTS

### ENGRAILED-1 is transferred from V1 interneurons to motoneurons in the ventral spinal cord

Previous studies reported *En1* expression in embryonic and early postnatal spinal cord V1 interneurons but, to our knowledge, no study has directly visualized EN1 in adult mouse V1 interneurons. The available studies identify *En1*-expressing cells in the adult mouse spinal cord using EN1-Cre-inducible reporter genes activated during development that maintain reporter gene expression throughout life (Sapir, 2004; Alvarez *et al*, 2005; Lane *et al*, 2021). In contrast with classical *in situ* hybridization (ISH) that gave no or very low signal in our hands, RNAscope ISH clearly reveals *En1* expression in adult 4.5-month-old mice V1 interneurons, including Renshaw cells (RCs) identified by *Calbindin-1* expression (Figure 1A). These *En1*-expressing cells are localized dorsally (V1^P^) and ventrally (V1^R^, RCs) to a population of *En1*-negative neurons identified by a choline acetyltransferase (*ChAT*) ISH (RNAscope) probe, thus MNs. *En1* expression, quantified by RT-qPCR, is fully maintained between 4.5 and 16 months and reduced by 2-fold in the *En1*-Het mouse (Figure 1B).

**Figure 1.**
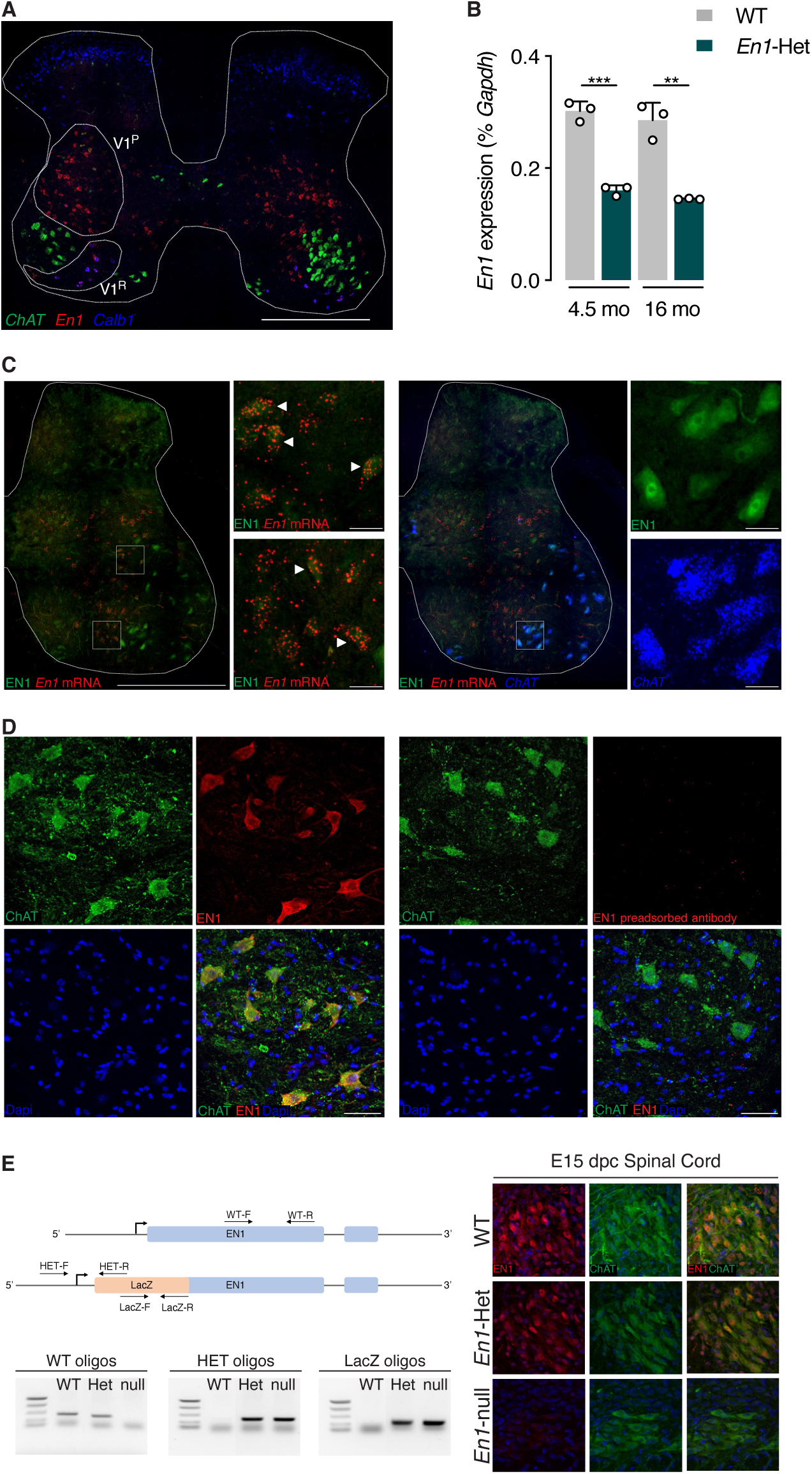
*En1* expression in the adult spinal cord. **A.** Triple RNAscope *in situ* hybridization showing *Engrailed-1* (*En1*), *Choline acetyltransferase* (*ChAT*) and *Calbindin-1* (*Calb1*) expression in the lumbar spinal cord. *En1* is expressed in V1 interneurons, dorsal (V1^P^) and ventral (V1^R^) to the main *ChAT*-expressing motoneuron pool. Ventral interneurons correspond to Renshaw cells as shown by *Calb1* expression. Scale bar: 500 μm. **A. B.** RT-qPCR of RNA from the lumbar enlargement at 4.5- and 16-months of age shows stable *En1* expression in WT at both ages and a two-fold reduction of expression in heterozygous mice. Unpaired two-sided t-test. **p<0.005; ***p<0.0005. n=3. **B.** Triple staining EN1 IHC (green), *En1* RNAscope ISH (red) and *ChAT* RNAscope (blue) demonstrating the double-staining of *En1* mRNA and protein (EN1) in the V1 interneuron population (left panel insets, arrowheads point towards examples of double-stained V1 interneurons), and the presence of EN1 protein in large cells not expressing *En1* mRNA (left panel) but expressing *ChAT* (right panel insets). Scale bar: 500 µm, 30 µm for high magnification insets. **C.** Left: EN1 (red) detected with the LSBio antibody is localized in ChAT-expressing neurons (green) in the ventral horns of the spinal cord. Right: EN1 signal is lost upon preincubation of the antibody with 1.5 M excess of recombinant hEN1. Scale bar: 50 µm. **D.** Left: Relative positions of the different oligonucleotides selected to genotype the E15 embryos and examples of the genotyping based on the combination of PCRs with the different pairs of primers. Right: Double ChAT/EN1 immunostaining demonstrating the co-localization of the two proteins in the WT and the *En1*-Het ventral cord and the absence of EN1 staining in MNs from *En1*-KO embryos. Note that the staining is reduced in *En1*-Het embryos, compared to WT embryos.

To examine *En1* expression at the protein level, we tested several antibodies and found that LS-B9070 from Lifespan Biosciences (LSBio) directed against EN1 amino acids 1 to 30 gives a clear and consistent staining of both V1 interneurons (Figure 1C, left panels) and of *ChAT*-expressing (*ChAT*+) MNs (Figure 1C, right panels). Antibody specificity was verified in several ways. First, in spinal cord and ventral midbrain extracts, the LSBio antibody and a previously characterized 86/8 antibody (Sonnier *et al*, 2007), recognize a major band that migrates at the same position as recombinant EN1 (Figure EV1A). Second, immunostaining of ChAT+ cells by the LSBio antibody (Figure 1D, left panel) is totally lost upon preincubation of the antibody with recombinant hEN1 (right panel). Third, in *En1*-KO mice at embryonic day 15 (the KO is embryonic lethal), the staining is entirely absent in ChAT+ neurons (Figure 1E) demonstrating, in addition to antibody specificity, that EN1 transfers also at embryonic stages. Finally, serial antibody dilutions demonstrate that the intensity of ChAT-expressing (ChAT+) MN staining is decreased in adult *En1*-Het mouse compared to WT (Figure EV1B).

The presence of EN1 protein but not mRNA in MNs suggests that EN1 produced by the interneurons is efficiently secreted and internalized by adult spinal MNs. As will be described below, EN1 intercellular transfer is further demonstrated by its extracellular neutralization through the expression of a secreted single chain antibody (scFvEN1) directed against EN1 (Wizenmann *et al*, 2009), resulting in an important reduction of MN staining by the LSBio antibody (Figure 4A).

### *Engrailed1-*heterozygote mice show alpha-motoneuron retrograde degeneration

Because *En1*-expressing mDA neurons degenerate in the *En1-*Het midbrain (Sonnier *et al*, 2007), we examined whether ventral spinal cord neuronal populations, in particular interneurons that express *En1*, also degenerate in the *En1*-Het mouse. We used RNAscope labeling to compare the number of *En1*-expressing neurons in the spinal cord ventral horns of *En1-*Het mice and their WT littermates at 4.5 months of age. Figure 2A shows no loss of V1 interneurons, including RCs (*Calbindin1*-expressing cells). Given the absence of a change in *En1* expression between 4.5 and 16 months quantified by RT-qPCR (Figure 1B), these results demonstrate that V1 interneuron survival is not affected by reduced *En1* expression in the *En1-*Het mouse mutant. We also quantified the total number of *ChAT*+ motoneurons in 4.5-month-old mice and did not observe a reduction in the *En1*-Het mouse (Figure 2A).

**Figure 2.**
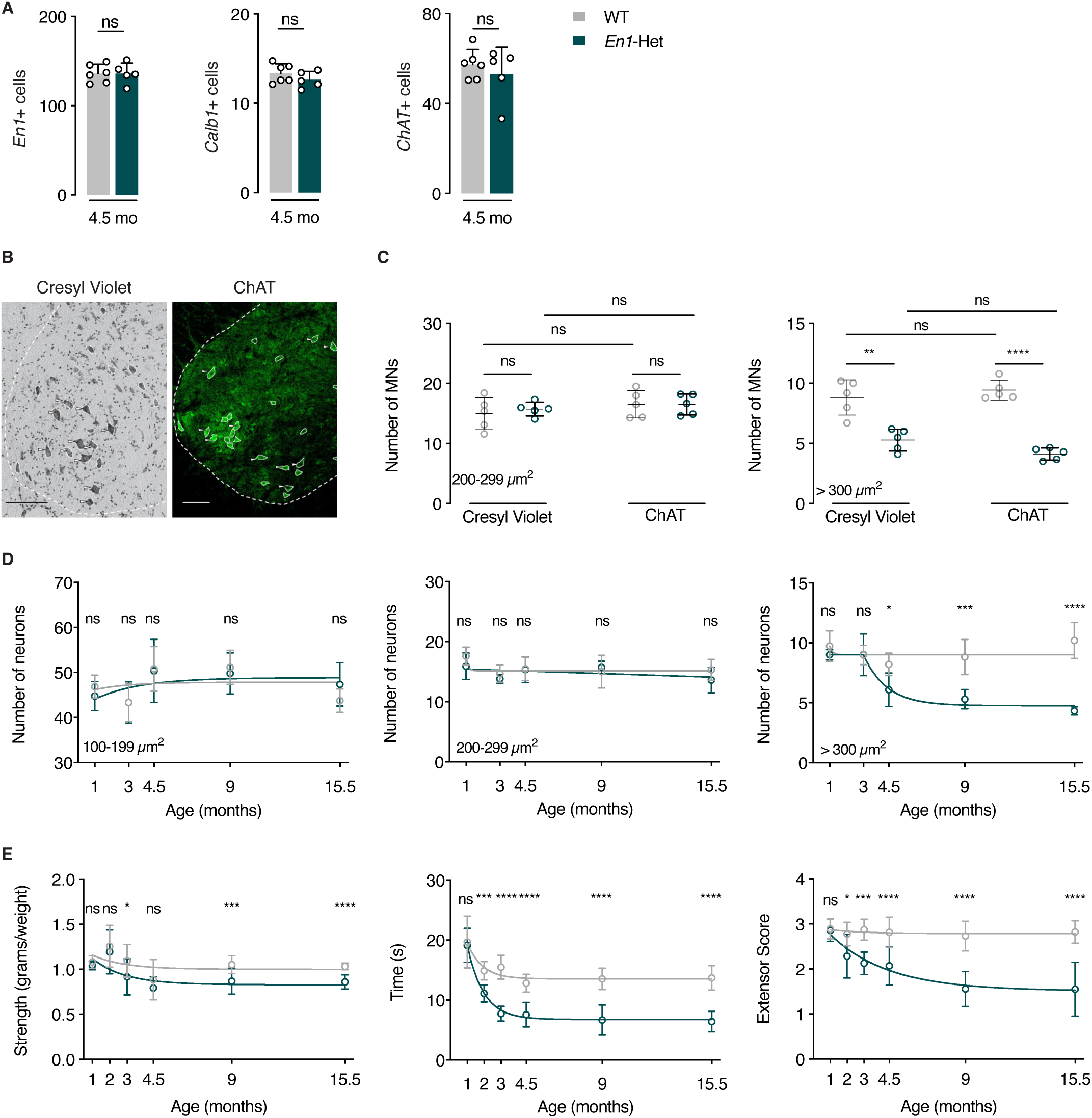
Progressive loss of αMNs and strength decrease in *En1*-HET mice. **A.** Analysis of the number of *En1*+, *Calb1*+ and *ChAT*+ neurons (triple RNAscope). At 4.5 months, WT and *En1*-Het mice show no difference in the number of cells expressing *En1*, *Calb1* or *ChAT*. n=5-6. **B.** Cresyl violet and ChAT staining of a ventral WT spinal cord at the lumbar level. Scale bar: 100 µm. **C.** Cresyl violet and ChAT staining at the lumbar level show no medium size (200-299 µm^2^) cell loss (γMNs) and a decrease of about 50% in the number of large size (>300 µm^2^) cells (αMNs). Unpaired two-sided t-test. **p<0.005; ***p<0.0005. n=5. **D.** Lumbar level Cresyl violet staining shows that, in contrast with small and medium size neurons (interneurons and γMNs, left and center panels), large size neurons (αMNs, right panel) undergo progressive death first measured at 4.5 months. The values represent the average number of cells per ventral horn. For the small neurons (100-199 µm^2^), there was no main effect, two-way ANOVA for repeated measures for treatment group: F(1,43)=0.0017, p=0.968, ns. For the medium sized neurons (200-299 µm^2^) two-way ANOVA for repeated measures for treatment group: showed no main effect F(1,43)=2.085, ns. For the large neurons (>300 µm^2^), two-way ANOVA for repeated measures showed a significant main effect F(1,43)=59.99, p<0.0001. Post-hoc comparisons were performed by unpaired two-sided t-test with equal SD comparing WT with *En1-*Het at each time point (*p<0.05; **p<0.005; ***p<0.0005; ****p<0.0001). n=5 to 6. **E.** Compared to WT mice, *En1*-Het mice experience gradual strength loss. This loss is observed with the forepaw grip strength (left panel), the inverted grid test (center panel and the hindlimb extensor reflex test (right panel). Strength loss is first observed between 2 and 3 months, thus before measurable αMN cell body loss. Two-way ANOVA showed significant main effects for grip strength (F(1,136)=19.18, p<0.0001), inverted grid test (F(1,103)=143.1, p<0.0001) and extensor score (F(1,103)=10.1, p<0.0001). Comparisons were made by unpaired two-sided t-test with equal SD comparing WT with *En1-*Het at each time point (*p<0.05; **p<0.005; ***p<0.0005; ****p<0.0001). n=4 to 20. The analysis of neuron number in A, the direct comparison of Cresyl violet and ChAT in C and the longitudinal studies in D and E were performed once.

The ChAT-expressing population is heterogeneous with a mixture of γMNs and αMNs and the global analysis of *ChAT+* cells in Figure 2A may have missed degeneration, total or partial, of one of the two populations. Gamma and αMNs differ by size with the surface of γMNs ranging from 200 to 300 µm^2^ and that of αMNs equal or greater than 300 µm^2^ (Powis & Gillingwater, 2016). We counted MNs on Cresyl violet-stained and ChAT-labeled lumbar enlargement spinal cord sections (Figure 2B), taking soma size into account. We first performed this analysis at 9 months, reasoning that any degeneration would be more apparent at later ages. Cresyl violet and ChAT staining yield identical results and demonstrate a specific loss of large (>300 µm^2^) neurons (Figure 2C). The concordance between data obtained from Cresyl violet and ChAT immunostaining precludes that the loss of ChAT+ cells in *En1*-Het mice is a consequence of ChAT expression down-regulation. We used Cresyl violet to quantify the number of small (100-199 µm^2^), medium (200-299 µm^2^) and large (>300 µm^2^) neuronal populations as a function of age. Figure 2D demonstrates that, in the *En1*-Het mutant, the number of small-size (100-199 µm^2^) and intermediate (200-299 µm^2^) neurons does not decrease with age (Figure 2D, left and middle panels). In contrast, the number of αMNs strongly decreases between 3 and 4.5 months of age, with about 50% loss at 9 months and no further decrease between 9 and 15.5 months (Figure 2D, right panel). The plateau in αMN death is best explained by the fact that, with time and due to αMN death, each remaining MN receives a higher amount of EN1 (Figure EV2) which may slow down neurodegeneration. We then determined the consequences of αMN degeneration on muscle strength by measuring forepaw grip strength, the time holding onto an inverted grid, and the hindlimb extensor reflex. Figure 2E shows that strength loss appears between 2 and 3 months and reaches a maximum between 4.5 and 9 months of age. The loss of strength between 2 and 3 months therefore precedes αMN loss first observable at 4.5 months (Figure 2D, right panel).

To further characterize αMN degeneration, we analyzed the organization of neuromuscular junctions (NMJs) by following the number of acetylcholine receptor (AChR) clusters, the endplate area, the percentage of endplates with perforations, and that of fully occupied endplates (Sleigh *et al*, 2014). The number of AChR clusters identified with fluorescent α-bungarotoxin (α-BTX) did not change with time (Figure 3A) and the endplate area remained stable for 9 months, with a modest 20 to 25% decrease at 15.5 months. A similar late-occurring (15.5 months) and modest effect was also observed for the percentage of endplates with perforations, a maturation index that is decreased in the mutant mice only at 15.5 months (Comley *et al*, 2016). In contrast, a stronger and earlier difference was observed in the percentage of fully occupied endplates identified by the binding of fluorescent α-BTX and immunostaining of the high molecular weight neurofilament (2H3) and the synaptic vesicle glycoprotein SV2A (Figure 3B, left panel). Fully occupied endplates were defined by an overlap between SV2A/2H3 and α-BTX staining (pixel analysis) occupying more than 80% of the endplate (see Methods). Using this criterion, significant retrograde degeneration could already be measured in 3-month-old animals (Figure 3B, right panel). Quantitative RT-PCR and EN1 immunohistochemistry failed to reveal EN1 protein or mRNA at the level of the endplate, precluding that degeneration is a consequence of EN1 partial loss of function at the endplate level (Figure EV3).

**Figure 3.**
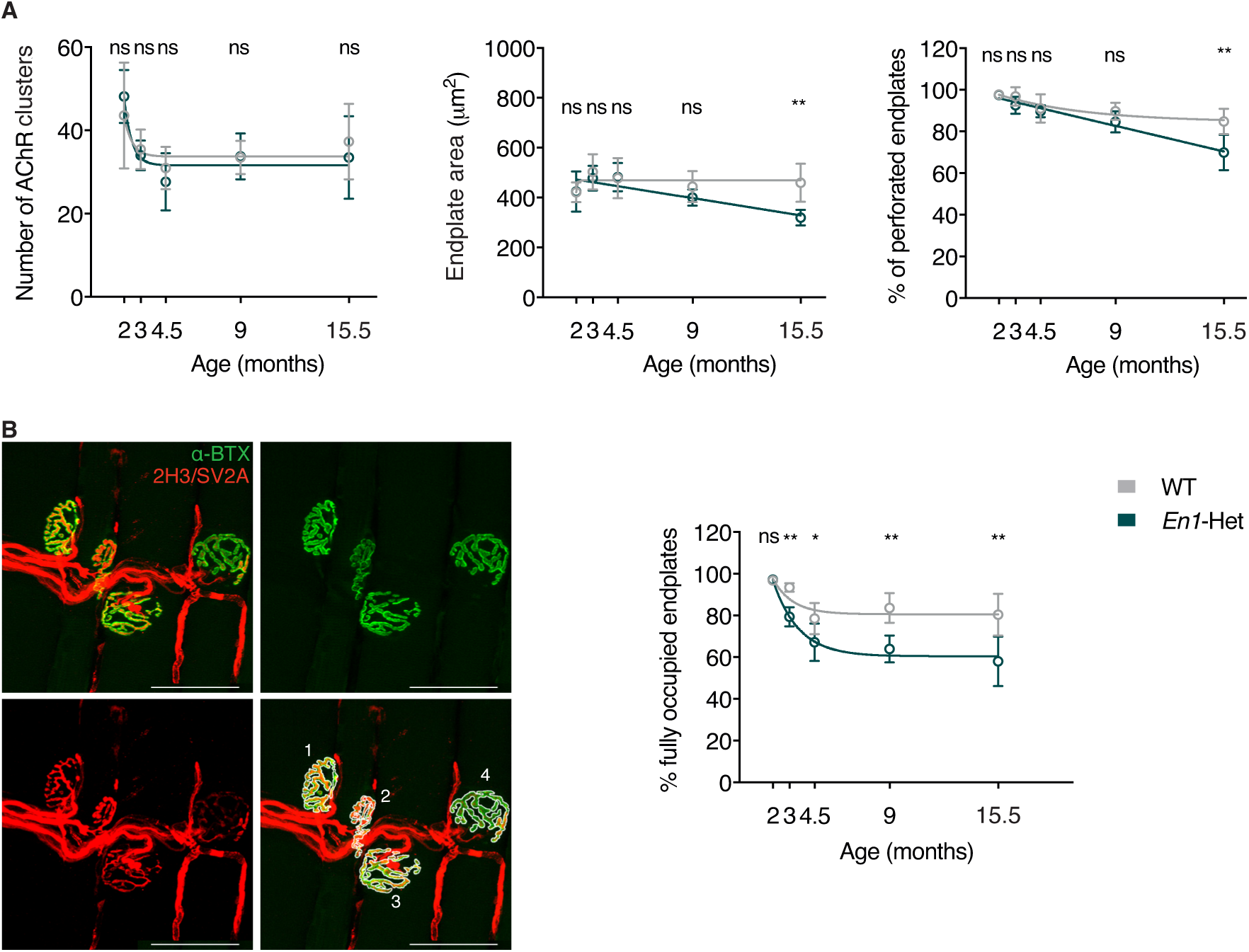
Neuromuscular junction (NMJ) morphology differences between WT and *En1-* Het mice. **A.** NMJs of *En1-*Het and WT mice show similar numbers of AChR clusters (left panel) and late-occurring (15.5 months) decrease in endplate area (center panel) and percentage of perforated endplates (right panel). The number of AChR clusters seems to decrease between 2 and 3 months, but two-way ANOVA showed no significant genotype effect (F(1,47)=0.1291, ns). There was a significant main effect for endplate area (F(1,47)=5.778, p=0.0202) and for the percentage of perforated endplates (F(1,47)=13.82, p=0.0005). Post hoc analysis revealed significant genotype differences at 15.5 month of age. Unpaired two-sided t-test with equal SD comparing WT with *En1-*Het at each time point (**p<0.005). n=4 to 8. **A. B.** Left panel illustrates the use of Alexa Fluor 488-conjugated α-bungarotoxin (α-BTX, in green) and of neurofilament and synaptic vesicle glycoprotein antibodies (2H3 and SV2A, in red) to evaluate the percentage of fully occupied endplates (>80% occupancy). The right panel shows that the % of fully occupied endplates decreases progressively in the *En1*-Het mouse, starting between 3 and 4.5 months of age. Scale bar: 50µm. Two-way ANOVA showed a significant main effect (F(1,47)=45.45, p<0.0001). Unpaired two-sided t-test with equal SD comparing WT with *En1-*Het at each time point (*p<0.05; **p<0.005). n=4 to 8. See details of analysis in the Methods section. The longitudinal study in A and B was performed once.

All in all, muscle weakness and NMJ changes preceding cell body loss demonstrate selective retrograde degeneration of αMNs in the *En1*-Het mouse.

### Extracellular neutralization of ENGRAILED-1 partially reproduces the *Engrailed1*-heterozygote phenotype

Since αMNs do not express *En1*, their degeneration in the *En1-het* mouse must be a consequence of the *En1* heterozygote status of the V1 interneurons with several mechanisms contributing, alone or in combination, to the phenotype. In light of the presence of EN1 in MNs (Figure 1), of its known direct non-cell autonomous activity in several developmental phenomena (Layalle *et al*, 2011; Wizenmann *et al*, 2009; Brunet *et al*, 2005; Rampon *et al*, 2015; Amblard *et al*, 2020b), and of EN1/2 protective activity on mDA neurons (Sonnier *et al*, 2007; Alvarez-Fischer *et al*, 2011; Blaudin de Thé *et al*, 2018; Rekaik *et al*, 2015; Thomasson *et al*, 2019), a parsimonious hypothesis is that EN1 exerts a direct non-cell autonomous trophic activity on αMNs.

To investigate this possibility, we used a strategy successfully developed in previous studies consisting in the expression of a secreted neutralizing anti-EN1 single-chain antibody (scFvEN1) (Wizenmann *et al*, 2009) in the spinal cord of WT mice. The antibody, or its inactive control harboring a single cysteine to serine mutation that prevents disulfide bond formation and EN1 recognition, were cloned into an AAV8 under the transcriptional control of the Glial Fibrillary Acidic Protein (GFAP) promoter (Figure 4A). This promoter directs scFvEN1 expression and secretion by astrocytes, thus allowing for the neutralization of EN1 in the extracellular space. Viruses were injected intrathecally (IT) at lumbar level 5 (L5) in 1-month-old WT mice, allowing for robust scFvEN1 expression one month later (Figure 4B). The amount of EN1 in MNs was quantified 6 months after virus injection, demonstrating a strong reduction in mice expressing scFvEN1 but not the inactive mutated form (Figure 4C).

**Figure 4.**
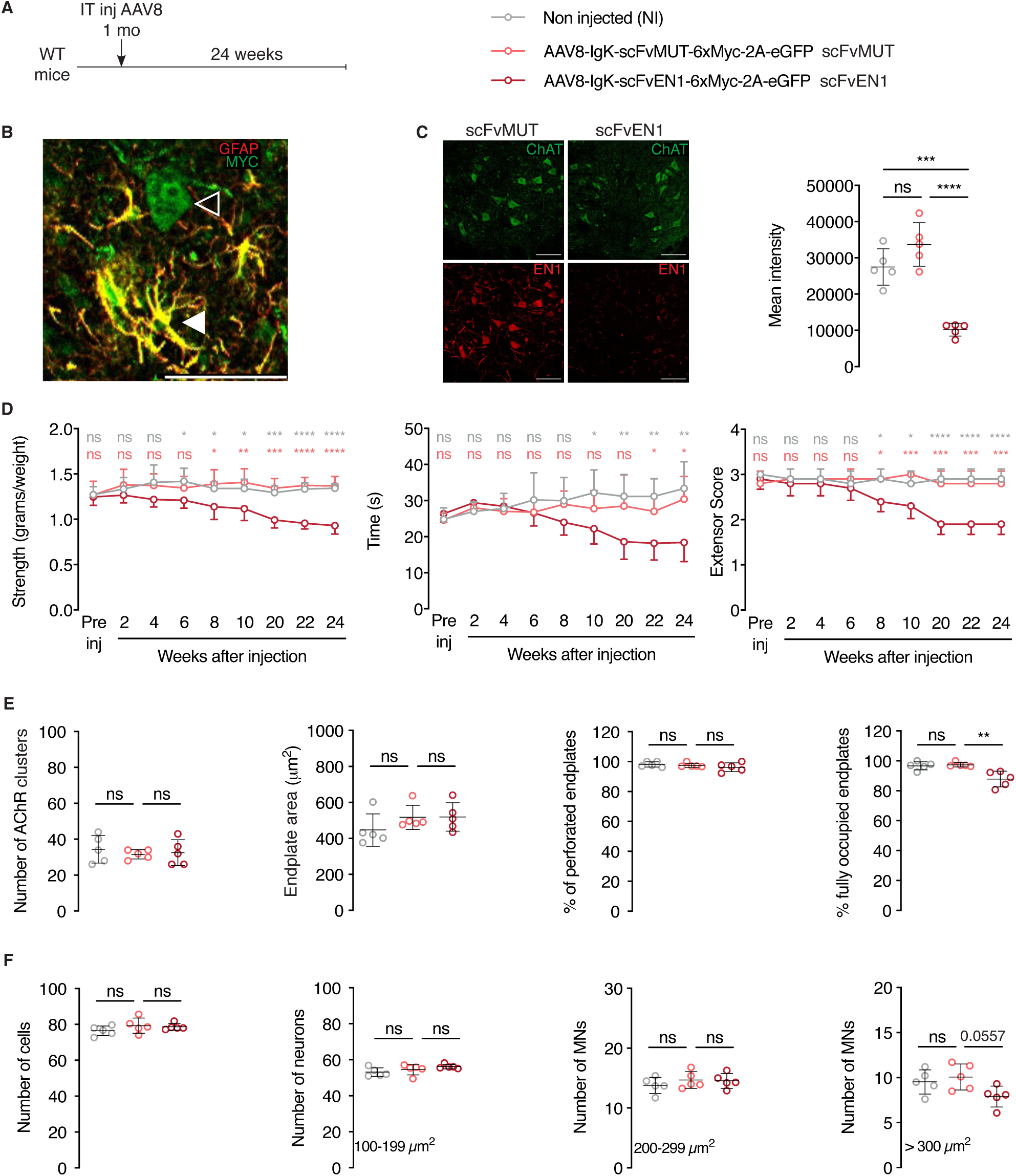
The effects of extracellular EN1 neutralization on strength, MN survival, and NMJ morphology. **A.** Experimental paradigm and structure of AAV8-encoded constructs containing glial fibrillary acidic protein (GFAP) promoter for expression in astrocytes, Immunoglobulin K (IgK) signal peptide for secretion, anti-ENGRAILED single-chain antibody (scFvEN1), 6 myc tags (6xMyc), skipping P2A peptide, and enhanced Green Fluorescent Protein (EGFP). An inactive control antibody (scFvMUT) contains a cysteine to serine mutation that prevents disulfide bond formation between IgG chains, thus epitope recognition. The AAV8 were injected in 1-month-old WT mice and the strength phenotypes were followed for 6 months before anatomical analysis. **B.** Analysis 1-month post-injection showing that scFv antibodies are expressed in astrocytes (white arrowhead) double-stained for GFAP and Myc and exported (empty arrowhead). Scale bar: 50 µm. **C.** Left panel illustrates that expressing the scFvEN1, but not scFvMUT, abolishes EN1 staining by LSBio anti-EN1 antibody in ventral horn ChAT+ cells and right panel quantifies this inhibition 1-way ANOVA followed by Tukey corrected post-hoc comparisons. (n=5 mice per group, ***p<0.0005, ****p<0.0001). Scale bar: 100 µm. **D.** The three graphs illustrate how the WT antibody but not its mutated version leads to progressive strength decrease. Two-way ANOVA showed significant main effects for grip strength (F(2,12)=15.88, p<0.0005), inverted grid (F(2,107)=19.86, p<0.0001), and extensor score (F(2,12)=30.22, p<0.0001) followed by Tukey corrected post-hoc comparisons. (*p<0.05; **p<0.005; ***p<0.0005; ****p<0.0001). n=5 per treatment. **E.** Six months following infection (7-month-old mice), extracellular EN1 neutralization does not modify the number of AChR clusters, nor the endplate surface area, nor the percentage of perforated endplates. In contrast, the percentage of fully occupied endplates is diminished (right end panel). 1-way ANOVA followed by Tukey corrected post-hoc comparisons. (**p<0.005). n=5 per treatment. **F.** Six months following infection, extracellular EN1 neutralization does not globally modify the total neuron number of cells at the lumbar level (left panel). A separate analysis of small (100-199 µm^2^), medium (200-299 µm^2^), and large (>300 µm^2^) neurons demonstrate a specific (p<0.0557) loss of the latter category (αMNs). 1-way ANOVA followed by Tukey corrected post-hoc comparisons. N=5 per treatment. The extracellular neutralization study was performed twice.

In the three strength tests (anterior limb grasping, holding onto an inverted grid and hindlimb extensor reflex), mice expressing extracellular scFvEN1 experience gradual strength loss that begins between 6 and 8 weeks after virus injection (Figure 4D). In contrast, the performance of mice expressing the mutated inactive antibody are indistinguishable from that of WT animals. Endplate morphology and αMN survival were evaluated 6 months after injection of the antibody-encoding viruses (at 7 months of age). The results demonstrate no effect of EN1 extracellular neutralization on the number of AChR clusters, endplate area, and percentage of endplates with perforations, but a modest effect on the percentage of fully occupied endplates (Figure 4E, right panel). The analysis of the total number of cells and, separately, of the number of small, medium and large size neurons shows a small but specific loss of αMNs (Figure 4F).

Qualitatively, the latter results are similar to those obtained with the *En1*-Het mice with a small decrease in the number of fully-occupied endplates and the specific loss of large size αMNs. However, it is of note that the magnitude of the changes at 7 months was less than in the *En1*-Het mouse, with a 15% loss in fully occupied endplates and a 25% reduction in the number of αMNs, compared to 30% and 50% in 7-month-old *En1*-Het, respectively (Figures 2 and 3). This comparison, illustrated in Figure EV4, shows that secreted EN1 participates in the *En1-*Het phenotype but may not fully explain its magnitude.

### Exogenous ENGRAILED-1 rescues the *Engrailed1*-heterozygote phenotype

We have previously demonstrated that EN1 administration in the ventral midbrain promotes mDA neuron survival in mouse and macaque PD models (Sonnier *et al*, 2007; Alvarez-Fischer *et al*, 2011; Rekaik *et al*, 2015; Blaudin de Thé *et al*, 2018; Thomasson *et al*, 2019). Since extracellular EN1 exerts a trophic activity on αMNs, we evaluated whether exogenous administration of EN1 could prevent αMN death in *En1*-Het mice. Human recombinant EN1 (hEN1) was intrathecally injected at the L5 level at a concentration of 1 µg in 5 µL in 3-month-old *En1*-Het mice (Figure 5A), a time when all αMNs are still present but muscle weakness is already noticeable, a consequence of retrograde degeneration initiated at the endplate (Figure 3). One and a half months later, the single EN1 injection had restored strength as measured by the 3 tests of anterior limb grasping, holding onto an inverted grid, and hindlimb extensor reflex (Figure 5A). Accordingly, the percentage of fully occupied endplates was found to be back to WT levels (Figure 5B, left panel), in parallel with the prevention of αMN death (Figure 5B, right panel). This last series of experiments establishes that extracellular EN1 exerts a curative trophic activity on αMNs of already symptomatic mice allowing for endplate reinnervation and the arrest of αMN degeneration.

**Figure 5.**
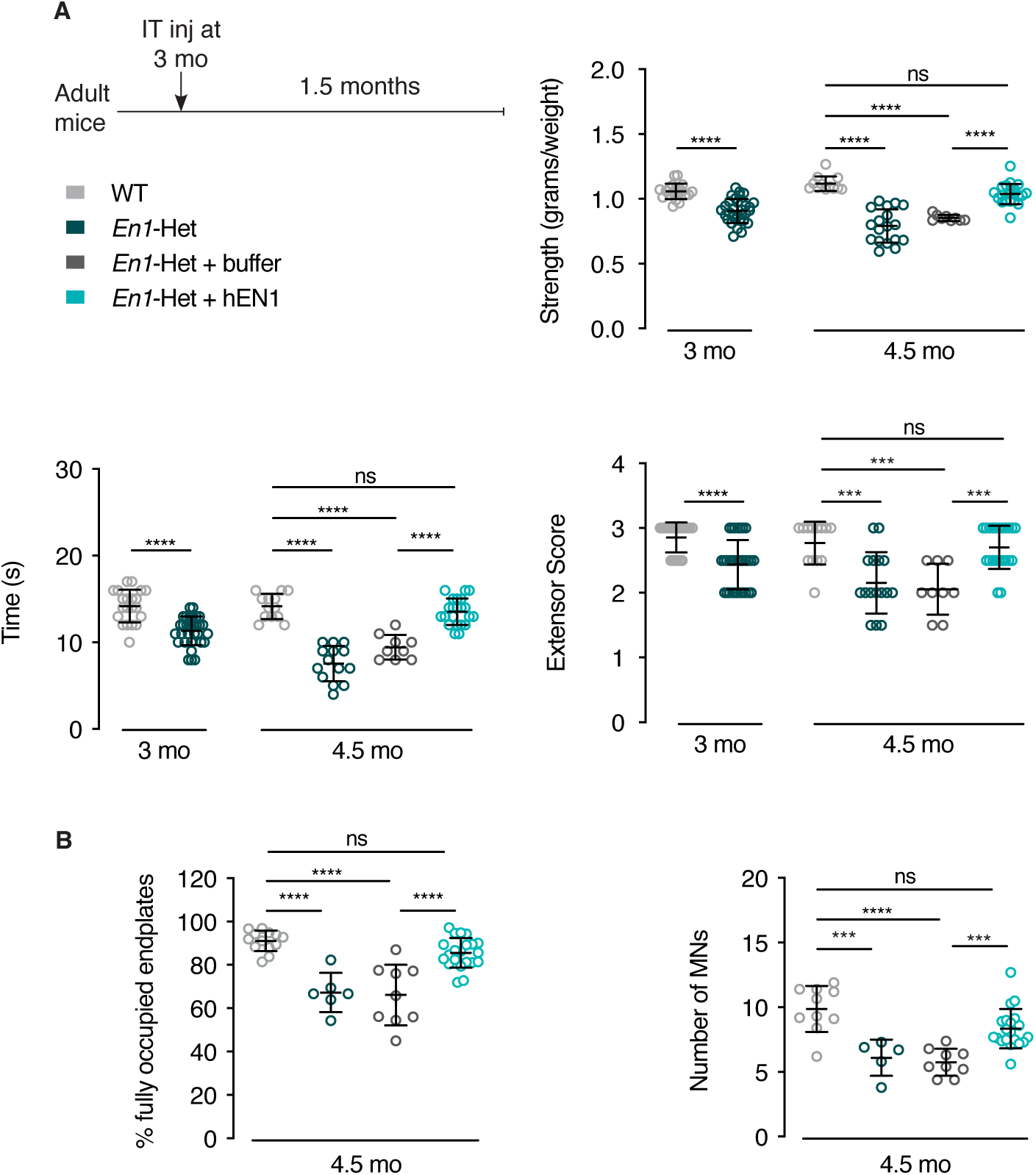
Intrathecal hEN1injection in *En1-Het* mice restores strength and prevents αMN death. **A.** Mice were tested for strength at three months of age, before the onset of αMN loss, but when strength has already decreased in *En1*-het mice as measured in the forepaw grip strength, inverted grid, and extensor reflex tests (left side of each graph). The next day, the *En1*-het mice were separated into two groups. One group received buffer and the other group recombinant hEN1 (1 µg in 5 µl), injected at the L5 level. One and a half months later (4.5 months of age) *En1*-het mice injected with hEN1 have recovered normal strength, in contrast with non-injected mice or mice injected with buffer. Unpaired two-sided t-test used at three months of age. ****p<0.0001. For 4.5-month comparisons, 1-way ANOVA followed by Tukey corrected post-hoc comparisons. ***p<0.0005; ****p<0.0001. n=9 to 31 **B.** At 4.5 months, following hEN1 injection at 3 months, the percentage of fully occupied endplates and the number of αMNs are not significantly different from control values. 1-way ANOVA followed by Tukey corrected post-hoc comparisons. ***p<0.0005; ****p<0.0001. n=5 to 21. The weakness and reversal with hEN1 and the neuroprotection were replicated in five independent experiments.

The duration of the effect of EN1 injection on strength was evaluated over time. Figure 6A summarizes an experimental design consisting of a single EN1 injection at 3 months. Strength analysis (anterior grip, holding time on the inverted grid and extensor reflex score) demonstrates that the effect of a single injection is maintained until approximately week 12 post-injection and decreases thereafter. In spite of this decrease, the strength of EN1-injected *En1*-Het mice is still superior to that of age-matched non-injected controls 24 weeks post-injection. Analysis of the % of fully occupied endplates and αMN number at 24 weeks confirms the long-lasting effect of a single injection. This led us to test whether a second injection 12 weeks after the first one could prolong the protective effect. Figure 6B summarizes this experimental protocol and illustrates that a second injection prolongs strength for another 8 to 12 weeks, depending on the test. Moreover, the percentage of fully occupied endplates and the number of αMNs at 24 weeks were identical to those observed in WT siblings.

**Figure 6.**
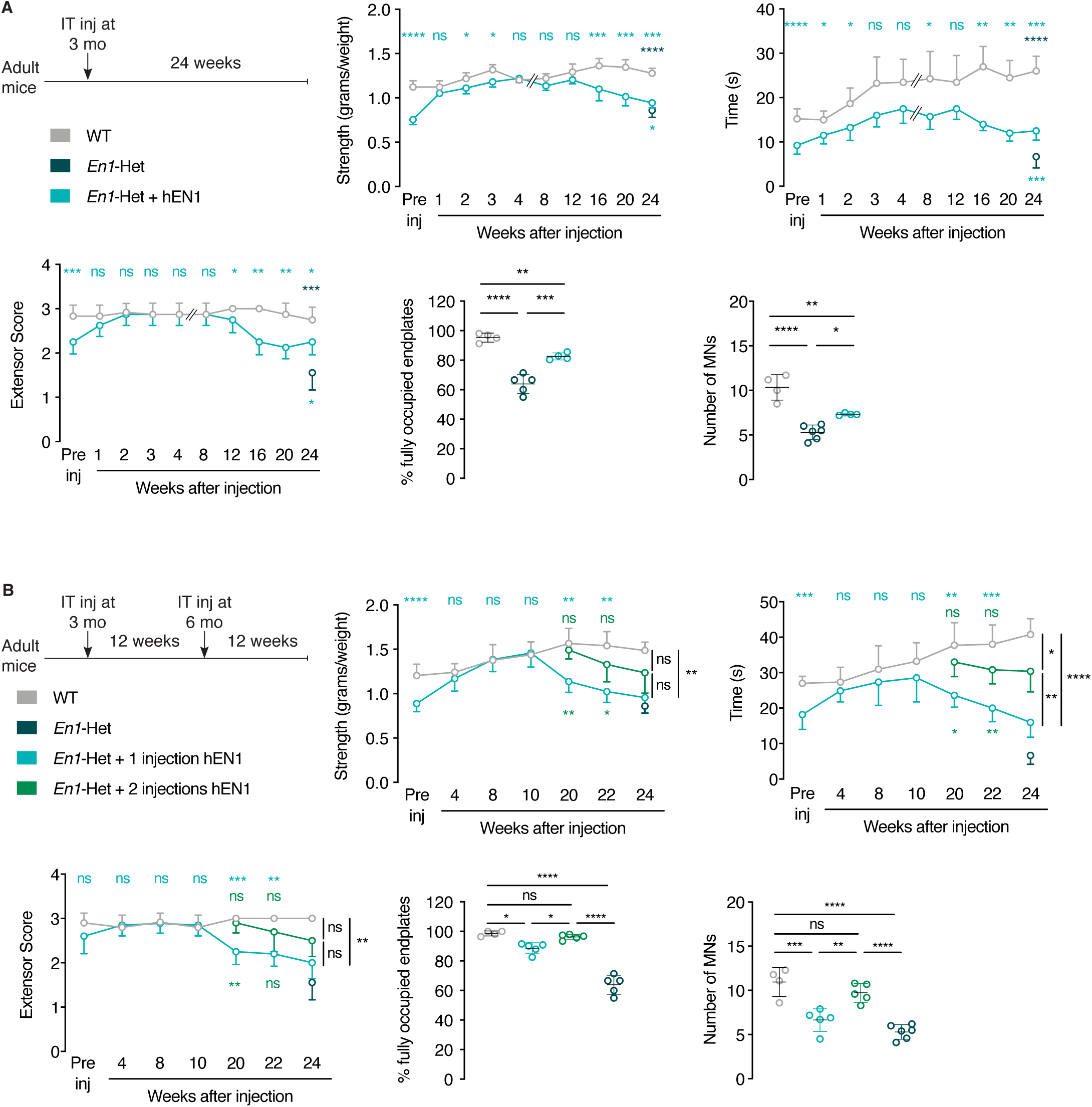
EN1 injection at 3 months has long-lasting effects on strength, endplate occupancy and αMN survival and the latter recoveries are prolonged by a second injection. **A.** Top left panel shows the single injection protocol whereby hEN1 is injected at 3 months (1µg in 5µl) intrathecally at the L5 level and mouse behavior followed for 24 weeks. The 3 time-course graphs demonstrate that a single injection restores strength measured by the tests of grip strength, time on the inverted grid and extensor reflex and that this effect lasts for 12 weeks. After 12 weeks strength decreases progressively but, even after 24 weeks, remains superior to that of untreated *En1*-Het mice. At 24 weeks, the % of fully occupied end-plates is inferior to that of WT mice, but superior to that of non-injected *En1*-Het mice. The same holds true for the number of αMNs. Two-way ANOVA revealed significant main effects for grip strength (F(1,76)=143.6, p<0.0001), inverted grid (F(1,76)=128.6, p<0.0001) and extensor score (F(1,76)=34.91, p<0.0001). At the different times the groups were compared by unpaired T-test with equal variances comparing WT with *En1-*Het injected at each time point. (*p<0.05; **p<0.005; ***p<0.0005; ****p<0.0001). For the endplate analysis and αMNs, groups were compared by 1-way ANOVA followed by Tukey corrected post-hoc comparisons (*p<0.05; **p<0.005; ***p<0.0005; ****p<0.0001). n= 4 to 10. **B.** Based on the results shown in A, a new experiment was performed with a second injection 12 weeks after the first injection at 3 months of age. Again, the injection at three months of age restored strength and in mice receiving the second injection strength was maintained an additional ten weeks or more compared to mice receiving a single injection. The three strength graphs demonstrate a positive effect of the second injection with values intermediate between those measured in WT mice and En1-Het mice with a single injection. The percentage of fully occupied endplates and the number of αMNs are back to wild type values in *En1*-Het mice injected twice. Two-way ANOVA revealed significant main effects for grip strength (F(2,81)=25.47, p<0.0001), inverted grid (F(2,91)=51.96, p<0.0001) and extensor score (F(2,104)=30.42, p<0.0001). At all times groups were compared by Unpaired T-test with equal variances through 12 weeks. After, the groups were compared by Tukey corrected post-hoc comparisons (*p<0.05; **p<0.005; ***p<0.0005; ****p<0.0001). For the endplate analysis and αMNs, groups were compared by 1-way ANOVA followed by Tukey corrected post hoc comparisons (*p<0.05; **p<0.005; ***p<0.0005; ****p<0.0001). n= 4 to 10. The single injection and two injection time course studies were performed once each.

### Intrathecally injected EN1 preferentially accumulates in MNs

We then followed EN1 localization after intrathecal injection and observed that 24 hours after injection EN1 is present in the perivascular space and in ventral horn cells, primarily in MNs characterized by ChAT expression (Figure 7A, upper row left panel). Figure 7A also illustrates that EN1 staining in MNs is transient, peaking between 6 and 24 hours, still visible at 48 hours and barely so at 72 hours. The latter images were taken with the 86/8 antibody that poorly stains endogenous EN1 allowing us to see primarily the exogenous protein, but similar images were obtained with the LSBio antibody. Motoneurons identified by ChAT-staining were separated by size to distinguish between αMNs and γMNs and the kinetics of internalization were quantified in the two subpopulations (Figure 7A, bottom row), using the 86/8 (left panels) and LSBio (right panels) antibodies. Indeed, the difference following internalization is much higher with the 86/8 antibody since the basal staining due to endogenous EN1 is very low, but the time courses obtained with the two antibodies are very similar. Furthermore, the data obtained with the LSBio antibody, which recognizes endogenous and exogenous EN1, suggest that exogenous EN1 at its peak (24 hours) doubles the concentration of EN1 per MN (assuming that the antibody recognizes equally endogenous and exogenous EN1). All in all, the latter series of experiments demonstrates similar patterns for αMNs and γMNs, with a very rapid accumulation that peaks at 24 hours and decays rapidly, reaching nearly basal levels at 72 hours, allowing one to estimate a short 24-hour EN1 half-life.

**Figure 7.**
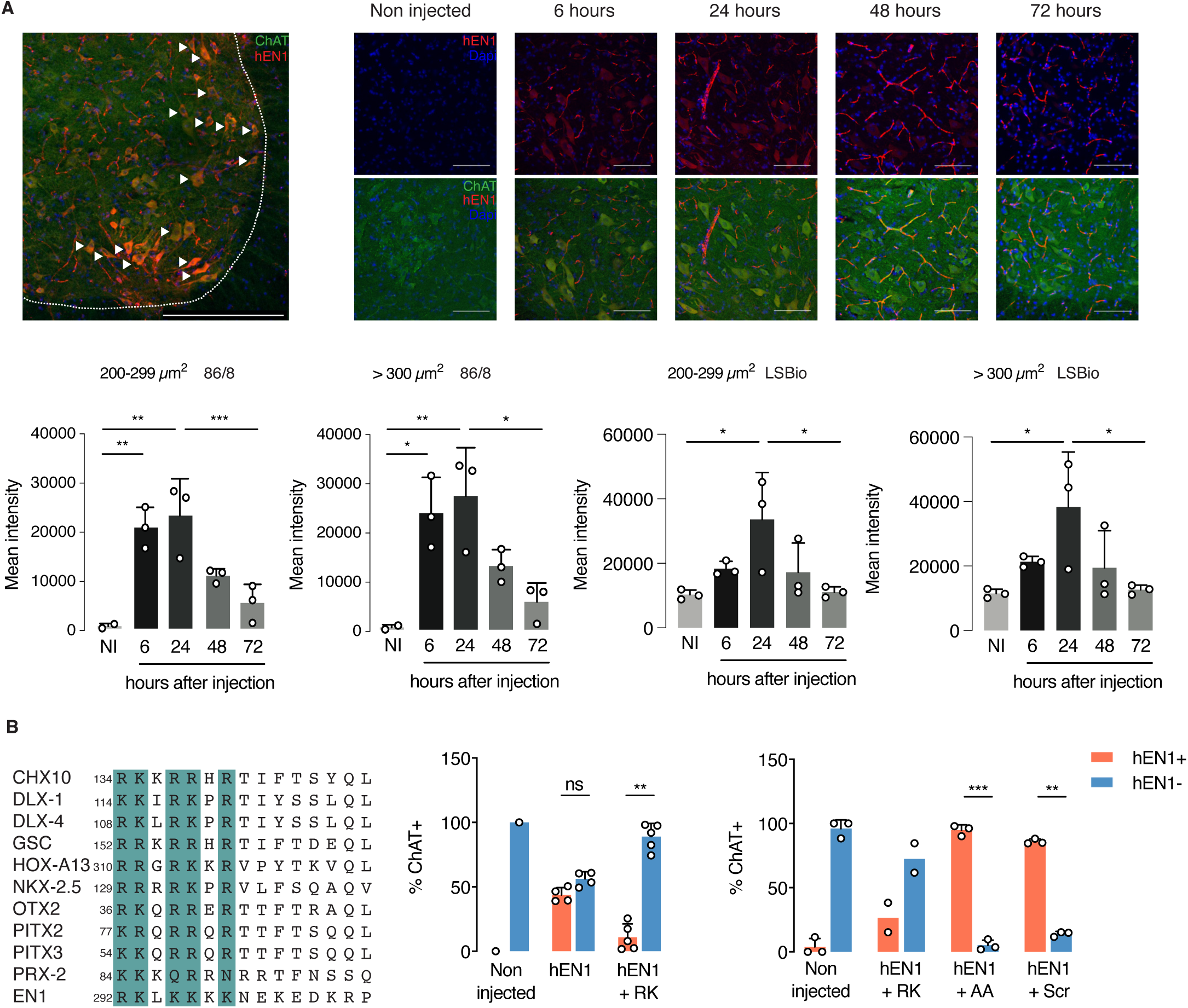
Following intrathecal injection, hEN1 is addressed specifically to all MNs. **A.** Top left panel shows that 24 hours after intrathecal injection (1 μg in 5µL) at the L5 level of 2-month-old mice, hEN1 (red) can be primarily visualized in ventral horn ChAT+ cells (green). Arrowheads show hEN1 internalized by MNs. Scale bar: 100 µm. Top right panels show the progressive accumulation and clearance of hEN1 in ventral horn MNs, with a peak between 6 and 24 hours. EN1 was revealed by the 86/8 antibody allowing for the visualization of exogenous EN1 only (scale bar: 100µm). Bottom panels show the quantification of EN1 in γMNs and αMNs with the 86/8 (two left panels) and the LSBio (two right panels) antibodies. Since the LSBio sees both endogenous and exogenous EN1, the increase is only 3-fold, but qualitatively, the results are very similar, demonstrating rapid internalization and clearance of the exogenous protein and allowing one to calculate a half-life of 24 hours. 1-way ANOVA followed by Tukey corrected post-hoc comparisons (*p<0.05; **p<0.005; ***p<0.0005). When not significant, p values are shown. n=3. The internalization experiments were performed once with each antibody. **B.** Left panel gives examples of putative glycosaminoglycan (GAG)-binding domain in 11 homeoprotein transcription factors. Based on the alignment and on published work on OTX2 (Beurdeley *et al*, 2012) and EN2 (Cardon *et al*, 2021), a putative EN1 GAG-binding domain (RK-EN1) was designed. Middle panel quantifies the inhibitory effect of RK-EN1 on hEN1 capture (86/8 antibody) by ChAT+ cells demonstrating that RK-EN1 in a 1 to 20 ratio reduces the % of EN1-postive MNs (ChAT+) from 50 to less that 10%. The right panel demonstrates that this inhibitory activity is not shared by the mutant AA peptide or by a scrambled (Scr) peptide. Unpaired two-sided t-test with equal SD (**p<0.005; ***p<0.0005). n=2-5. The GAG competition experiment was performed twice as described in the text.

The preferential capture of EN1 by MNs might reflect the presence of EN1 binding sites at the MN surface. We previously identified a glycosaminoglycan (GAG) binding domain within the homeoprotein OTX2 that allows for its preferential capture by cerebral cortex parvalbumin (PV) neurons (Beurdeley *et al*, 2012). This RK-OTX2 domain peptide (RKQRRERTTFTRAQL) competes for OTX2 binding to PV cells and antagonizes its specific internalization upon infusion or injection into the cerebral cortex^37^. A similar GAG-binding domain was recently described in EN2 (Cardon *et al*, 2023) and sequence comparisons between OTX2, EN2 and several other HPs identified putative GAG-binding sites in many of them (examples in Figure 7B). In the case of EN1, this domain (RKLKKKKNEKEDKRPRTAF), thereafter RK-EN1, is present between residues 292 and 310 of hEN1.

To test its potential role in EN1 localization, RK-EN1 was co-injected with EN1 (in a 20:1 molar ratio) to competitively inhibit EN1 access to MNs. Twenty-four hours after injection, EN1 gains access primarily to MNs, but the percentage of MNs that capture the protein is variable. In Figure 7B (central panel), EN1 gained access to 50% of all ventral horn MNs, characterized by ChAT expression, and this percentage dropped to 10% in presence of the competing RK-peptide, with most ChAT+ MNs devoid of EN1. In a separate experiment (Figure 7B, right panel), we tested peptide specificity by co-injecting EN1 with a peptide (AKLKAKKNEKEDKAPRTAF) in which one R and one K were replaced by alanine residues (AA peptide) or with a scrambled (Scr) RK-peptide (RLKNKEKPRDREKTKAKFK) composed of the same amino acids, but in a different order. For control of RK-EN1 activity, 2 mice were injected with EN1 and the RK-EN1 peptide, reproducing the inhibition of internalization observed in the first experiment (central panel). In contrast, the two variant peptides were devoid of inhibitory activity. The absence of inhibition by the scrambled peptide demonstrates that inhibition cannot be explained only by a positive charge effect, suggesting that RK-EN1 is an addressing sequence for MNs in the ventral mouse spinal cord. It is of note that, even though only αMNs degenerate in the *En1*-Het mouse, EN1 is specifically captured by both γMNs and αMNs.

### Bioinformatic analyses reveal p62/SQTSM1 as a regulated ENGRAILED-1 non-cell autonomous target in motoneurons

Retrograde degeneration of αMNs in the *En1*-Het and their long-lasting (2 months at least) rescue after a single EN1 injection is reminiscent of the situation described for mDA neurons, with the difference that EN1 activity is non-cell autonomous for αMNs and cell autonomous for mDA neurons (Rekaik *et al*, 2015). This similar protective effect led to investigate if genetic pathways interacting with *En1* might be common to αMNs and mDA neurons. To that end, we compared previously acquired RNA sequencing data of the SNpc in WT and *En1-*Het mice at 6 weeks of age, thus before the initiation of mDA neuron death but after the initiation of retrograde degeneration (Sonnier *et al*, 2007; Nordström *et al*, 2015). A significance threshold of p=0.05 was used, generating a list of differentially expressed genes (Rekaik *et al*, 2015) that was crossed with a library of MN-expressed genes (Bandyopadhyay *et al*, 2013). Several physiological pathways were selected (see Methods), resulting in a list of 402 genes. As in PD, there is ample evidence that the first steps in ALS occur in the distal motor neuron axon that retracts from the muscle in a ‘‘dying back’’ phenomenon (Fischer *et al*, 2004). To focus on relevant genes in the context of αMN retrograde degeneration, we used the STRING database to extract within the 402 genes those independently interacting with *SOD1*, *FUS*, *TARDBP-43* and *C9ORF72*, 4 genes mutated in a large majority of familial ALS (fALS). This resulted in a short list of 20 genes modified in mDA neurons following *En1* loss of function, expressed in MNs and in interaction with one or several of the 4 fALS genes (Figure 8A).

**Figure 8.**
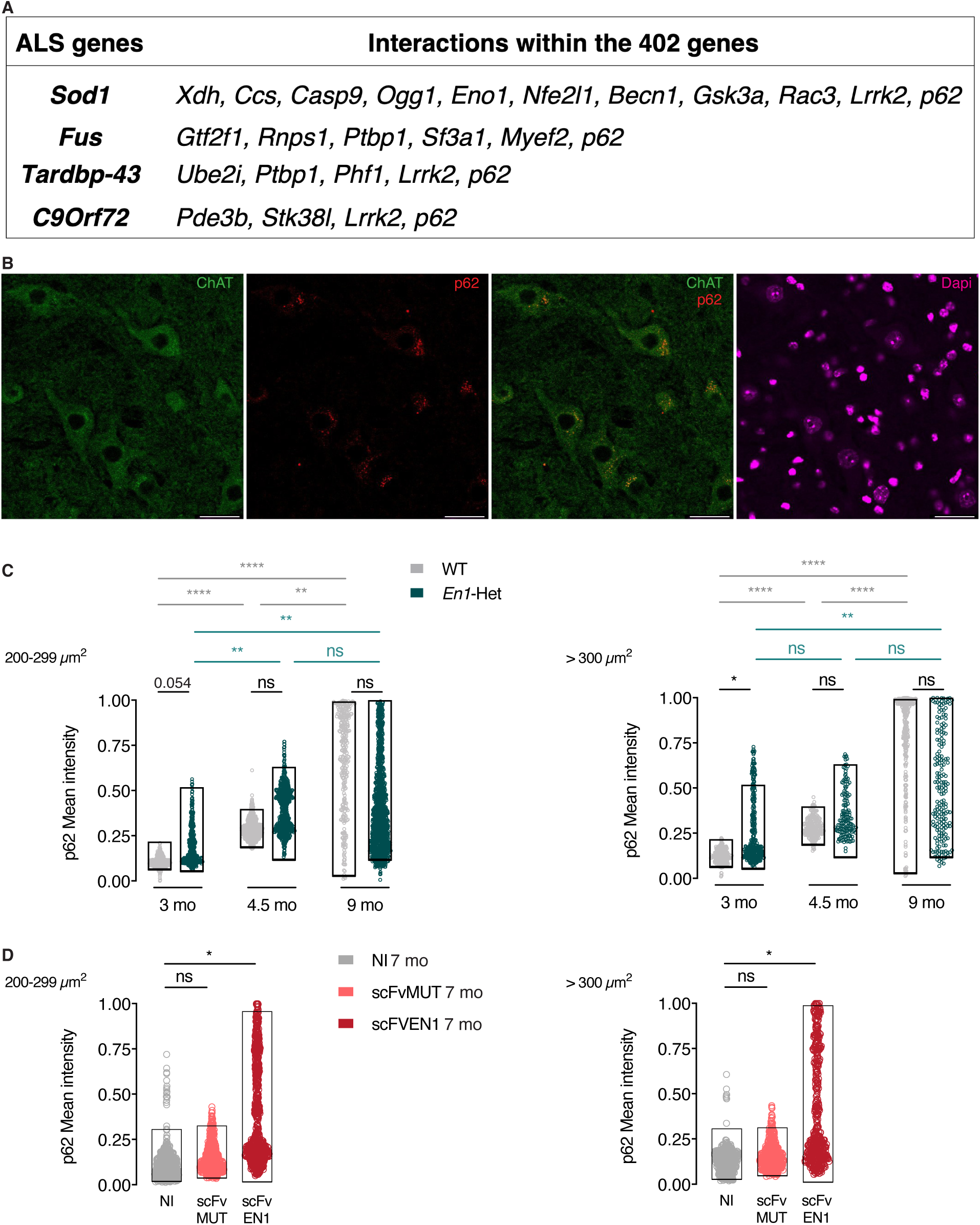
SQTSM1/p62 expression in MNs increases with age. **A.** The search for genes differentially expressed in WT and *En1*-Het mDA neurons and expressed in MNs, thus putative non-cell autonomous EN1 targets in MNs, allowed for the identification of 402 genes (after pathway selection). These genes were investigated for an interaction with genes mutated in the main 4 familial ALS forms. Among them, *p62/SQTSM1* (*p62*) expression is upregulated in the SNpc (RNA-seq) and in MNs of *En1*-Het mice. **B.** Immunohistochemical staining shows the presence of high amounts of p62/SQTSM1 in ChAT+ cell bodies of 3-month-old mice. Scale bar: 50 µm. **C.** Intensity measurements demonstrates that the mean of SQTSM1/p62 expression increases with age in WT γMNs (left) and αMNs (right). However, comparing WT and *En1*-Het shows a significant difference only at 3 months and not later. Statistical analysis is described under Methods. Between 151 and 1215 neurons and 3 to 5 mice were analyzed for each condition. **D.** Mean intensity of SQTSM1/p62 expression is increased in γMNs (left) and αMNs (right) in mice expressing scFvEN1 6 months after virus injection (7-month-old mice) demonstrating that EN1 extracellular neutralization increases SQTSM1/p62 expression. Expression of the mutated antibody (scFvMUT) does not increase p62 expression. Statistical analysis is described under Methods. Between 448 and 759 neurons and 5 mice were analyzed for each condition. The experiments in C and D were performed once each.

Among these 20 genes, *p62 or p62/SQTSM1* interacts with the four fALS genes, is mutated in sporadic forms of the disease (Shimizu *et al*, 2013; Chen *et al*, 2014; Fecto *et al*, 2011; Yang *et al*, 2015), and was recently shown to have variants in 486 fALS patients (Yilmaz *et al*, 2019). p62/SQTSM1 is a multifunctional protein that regulates the degradation of ubiquitinated proteins by the proteasome and carries ubiquitinated cargoes to the LC3 receptor at the autophagosome surface, a key step in cargo engulfment and lysosomal hydrolysis (Doherty & Baehrecke, 2018). The importance of protein degradation and autophagy in protein homeostasis and age-associated pathologies (Menzies *et al*, 2015; Leidal *et al*, 2018; Klionsky *et al*, 2021) led us to compare the immunostaining for p62/SQTSM1 in lumbar αMNs (>300µm^2^) and γMNs (200-299 µm^2^) in different conditions. A first observation (Figure 8B) is that expression of p62/SQTSM1 in WT ventral spinal cord is very strong in MNs characterized by ChAT expression. A second one is that the intensity of p62/SQTSM1 expression increases with age in WT αMNs and γMNs and may thus be considered as a MN physiological age marker (Figure 8C).

Comparison of p62/SQTSM1 expression in WT and *En1*-Het mice (Figure 8C), reveals a statistically significant or close to significant difference at 3 months in αMNs and γMNs (p=0.0499 and p=0.0536, respectively), before αMNs begin to degenerate, even though strength loss is already measurable (Figure 2C, D). Taking p62/SQTSM1 as an age marker, its increased expression at 3 months suggests accelerated ageing of αMNs and γMNs in *En1*-Het mice during this period, thus before αMN death and the ensuing reestablishment, at 4.5 and 9 months of age, of nearly normal EN1 concentration per MN (Figure EV2B). This led us to follow p62/SQTSM1 expression after extracellular EN1 neutralization in WT mice. Figure 8D shows that p62/SQTSM1 is significantly upregulated 6 months after injection of the AAV8-scFvEN1 in one-month-old mice, but not of its mutated inactive variant. It is of note that, due to a ceiling effect in measurements at 9 months, we used a ß distribution adjusted between 0 and 1 for all calculations in Figures 8C and 8D (see Methods).

At 3 months of age, all MNs (αMNs and γMNs) are still present in the *En1*-Het mouse but the enhanced p62/SQTSM1 expression suggests that mutant mice experience accelerated aging. We thus investigated if hEN1 injection at 1 month could antagonize MN premature aging at 3 months. Figure 9 describes the injection protocols and demonstrates that hEN1 injection prevents strength loss normally observed at 3 months using the three strength measurements (anterior limb grip strength, holding time on the inverted grid an extensor reflex). Figure 9B illustrates and quantifies p62/SQTSM1 expression, demonstrating that hEN1 intrathecal injection at 1-month antagonizes p62/SQTSM1 increase taking place at 3 months of age.

**Figure 9.**
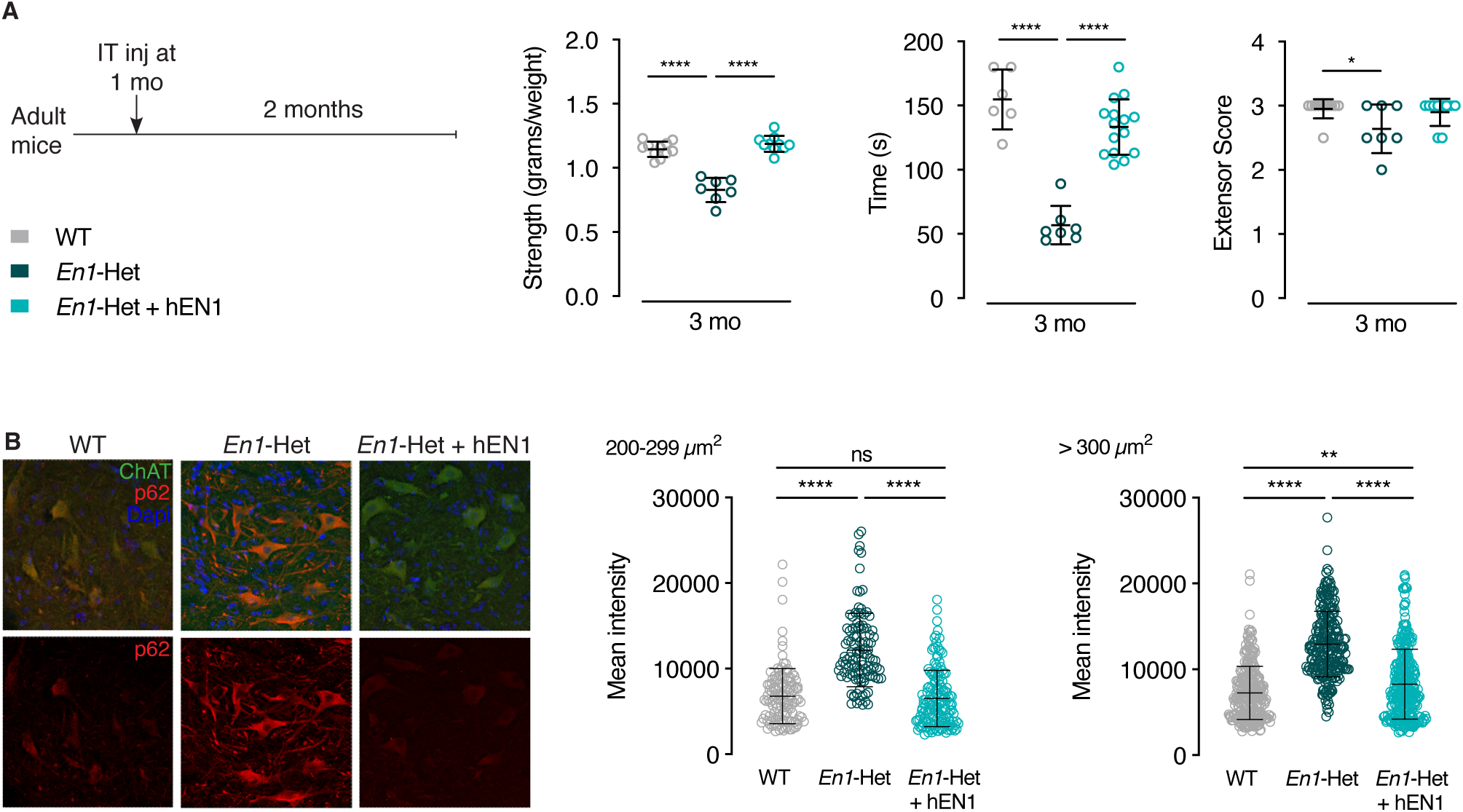
Human recombinant EN1 injection at 1 month prevents p62/SQTSM1 overexpression and strength loss in 3-month-old *En1*-Het mice. **A.** Left panel: Injection and analysis protocol. Right panel: muscle strength analysis demonstrating that hEN1 injection at 1 month prevents muscular strength decrease observed in 3-month-old En1-Het mice. **B.** Left panel: Increased p62/SQTSM1 staining in 3-month-old ChAT+ MNs from control *En1*-Het is abolished by hEN1 injection at 1 month. Right panel: quantification of p62/SQTSM1 staining in γMNs and αMNs of control and hEN1-injected 3-month-old *En1*-het mice. The behavioral analysis in A and the mean intensity analyses in B were performed once each.

Together, these results suggest that p62/SQTSM1 expression is a MN ageing marker, that the loss of an *En1* allele or EN1 neutralization in adult WT animals accelerate MN ageing and that hEN1 injection at 1 month of age prevents accelerated ageing and strength loss observed in 3-month-old *En1*-Het mice.

## DISCUSSION

This study establishes a novel adult function of EN1 and the importance of its expression in adult spinal cord V1 interneurons for αMN survival and muscle strength. This non-cell autonomous EN1 activity involves, at least in part, EN1 secretion as shown by the deleterious effect on the neuromuscular phenotype of extracellular EN1 neutralization. Moreover, hEN1 injected intrathecally accumulates in MNs. In the *En1*-Het mouse, this extracellular gain of function, following a single intrathecal injection of hEN1, antagonizes premature aging, reverses muscle denervation, arrests αMN degeneration, and restores normal strength for several weeks. While EN1 injections were done using the human recombinant protein, human and mouse EN1 are 91.1% identical and show similar efficacy in mDA neuron survival experiments (Blaudin de Thé *et al*, 2018; Rekaik *et al*, 2015; Alvarez-Fischer *et al*, 2011; Thomasson *et al*, 2019).

One day after injection, hEN1 is present around vascular structures probably in the perivascular space as well as within the parenchyma where it is observed within cells. Indeed, it has been shown previously that EN1 and EN2, similarly to many HPs(Lee *et al*, 2019), translocate across the plasma membrane and gain access to the cell cytoplasm and nucleus by a mechanism distinct from endocytosis (Prochiantz & Di Nardo, 2015; Di Nardo *et al*, 2018; Brunet *et al*, 2007; Amblard *et al*, 2020a; Di Nardo *et al*, 2020). Injected hEN1 shows preferential accumulation in ChAT-positive ventral MNs, although this does not preclude that some EN1 is present in the extracellular space and that a minority of other cell types may also capture the protein, albeit at lower levels, as suggested in Figure 7. The deleterious effect of blocking extracellular EN1 with an anti-EN1 scFv expressed in, and secreted by, astrocytes establishes that EN1 secreted by the V1 interneurons is internalized by αMNs and γMNs expressing EN1-binding sites and participates in several aspects of MN physiology, including αMN survival.

The latter finding is in line with the non-cell autonomous functions of EN1/2 reported in the fly wing disk and in the frog, fish, and chick optic tectum (Layalle *et al*, 2011; Wizenmann *et al*, 2009; Brunet *et al*, 2005; Rampon *et al*, 2015; Amblard *et al*, 2020b). It is also reminiscent of OTX2, another HP that specifically accumulates in PV interneurons of the cerebral cortex (Sugiyama *et al*, 2008). Prior to internalization, OTX2 binds to GAGs present at the surface of these interneurons thanks to a GAG-binding domain (RKQRRERTTFTRAQL) that overlaps with the first helix of the homeodomain (Beurdeley *et al*, 2012; Miyata *et al*, 2012). Interestingly, similar putative GAG-binding domains are present upstream of the homeodomain in many HPs (examples in Figure 7), including EN2 (Cardon *et al*, 2023) and EN1 for which this putative GAG-binding sequence is RKLKKKKNEKEDKRP. The role of this domain in the specific targeting of EN1 and the presence of EN1 binding sites at the MN surface are both illustrated by the ability of the peptide to abolish EN1 specific uptake by ventral spinal cord MNs. Although we have not directly demonstrated that EN1 binds GAGs, the conservation of the GAG binding domain and the analogy with OTX2 (Beurdeley *et al*, 2012) and EN2 (Cardon *et al*, 2021) support a specific association of EN1 with MN-expressed GAGs. If this is confirmed by future studies, it will be important to identify the molecular nature of this GAG as done for OTX2, EN2 and also VAX1 (Kim *et al*, 2014). Given the existence of a putative GAG-binding domain in many HPs, this study and those on OTX2 and VAX1 raise the issue of the existence of a sugar code for the specific recognition of their target cells by transferring HPs.

In the *En1-*Het mouse, deficits in strength begin before 3 months of age and might be a consequence of accelerated MN ageing, as suggested by the enhanced p62 expression at 3 months of age in mutant mice (discussed below). They coincide with a decrease in the percentage of fully occupied endplates first observed at 3 months and precede αMN loss which is initiated between 3 and 4.5 months. This temporal delay likely reflects a retrograde degeneration of αMNs that starts at the terminals, whereby peripheral strength is affected before αMN cell body loss. Retrograde degeneration also occurs in *En1*-expressing mDA neurons of *En1-*Het mice, with their terminals showing signs of degeneration at 4 weeks, whereas they only start dying at 6 weeks postnatal (Nordström *et al*, 2015). Future studies will be necessary to elucidate the molecular mechanisms that operate at the presynaptic and postsynaptic sites and might be in part regulated by EN1 transfer into MNs.

Retrograde degeneration of αMNs is also a prominent feature of human motoneuron diseases(Fischer *et al*, 2004). In this context, it is of note that hEN1 injection at 3 months not only prevents lumbar αMN death for months, but also restores normal endplate innervation and muscle strength, suggesting reinnervation. The neurodegenerative process is slowed or halted on a time scale of months and a second injection 3 months after the first one prolongs rescue for another 2 to 3 months. The time course of αMN degeneration in *En1*-Het animals is rapid with approximately 70% of final αMNs death taking place between months 3 and 4.5. Then, the process slows down to reach a plateau with ∼40% of the αMNs remaining at 15.5 months. It is thus possible that αMNs are heterogeneous in terms of their dependency on EN1. A more likely possibility is that when αMNs death gets close to 50%, the survivors receive sufficient secreted EN1 to halt accelerated ageing and the ensuing degeneration. This hypothesis is supported by Figure EV2B and the fact that the difference in p62/SQTSM1 expression is only visible at 3 months when MNs receive half of the normal EN1 dose but not at 4.5 months or later when almost half of the αMNs have disappeared. A strict evaluation of αMN response to decreasing EN1 doses will be necessary to fully explore this hypothesis.

Although EN1 activity is cell autonomous in mDA neurons and non-cell autonomous in αMNs, the similarities of their responses to *En1* hypomorphism, such as progressive retrograde degeneration preceding cell body loss, led us to interrogate the repertoire of differentially expressed genes in WT and *En1*-Het mDA neurons, and to compare it with the MN transcriptome. This produced a short list of genes and one of them, *p62*/*SQSTM1*, interacts with *SOD1*, *FUS*, *TARDBP-43* and *C9ORF72*, 4 genes responsible for a majority of fALS. p62/*SQSTM1* encodes p62/SQTSM1, a known regulator of degradation by the proteasome and autophagy through its role in bringing ubiquitinated proteins to the proteasome or to LC3 at the autophagosome surface (Klionsky *et al*, 2021). This unbiased identification of *p62*/*SQSTM1* in the *En1* “pathway” is particularly interesting because, in addition to interacting with the 4 main fALS mutations, it presents variants in 486 patients with familial ALS and is also mutated in sporadic ALS cases (Shimizu *et al*, 2013; Fecto *et al*, 2011; Chen *et al*, 2014; Yang *et al*, 2015). As already mentioned, p62/SQTSM1 expression in WT mice increases during ageing and can be taken as an age-marker. Compared to WT, we observe an upregulation of p62/SQTSM1 in the *En1*-Het mouse at 3 months suggesting accelerated aging. This accelerated aging, antagonized by hEN1 injection at 1 month, is best explained, in absence of αMN death at this age, by a dilution of EN1 content per MN. Similar upregulation, suggesting accelerated aging, is also measured in the scFvEN1 injected WT mice at 7 months when αMN death is limited to 20%, resulting in a significant decrease in EN1 content per MN (Figure 4C).

These series of observations can be interpreted in two non-mutually exclusive hypotheses. A first one is that p62/SQTSM1 plays a homeostatic role and that its increase represents a reaction against degeneration in ageing neurons. A second, based on the fact that p62/SQTSM1 itself is degraded through autophagosomes, is that ageing reduces autophagy and fosters p62/SQTSM1 accumulation (Bjørkøy *et al*, 2009; Jakobi *et al*, 2020; Jaakkola & Pursiheimo, 2009; Tai *et al*, 2016; Sun *et al*, 2023). P62/SQSTM1 is a multifunctional protein with distinct domains interacting with different co-factors (Ma *et al*, 2019). Thus, if a protective activity is demonstrated in further studies, specific deletions will be necessary to identify the pathways involved, in particular within the N-terminal domain, which presents a pro-survival activity through the N-kB pathway, and the C-terminal domain, which regulates proteostasis through the activation of proteasome and autophagosome activities (Foster & Rea, 2020).

Although EN1 is internalized by all MNs, which show similar increases in p62/SQTSM1 expression in both the *En1*-Het mouse or following EN1 extracellular neutralization, only αMNs and not γMNs degenerate in both models. This demonstrates that γMNs, in contrast with αMNs, are EN1-independent for their survival. The survival of approximately half of the αMNs may reflect a differential sensitivity of αMN subpopulations and/or the fact that, as discussed above, the surviving neurons receive higher and sufficient amounts of EN1. The absence of dependency on EN1 for γMN survival also applies to *En1*-expressing V1 interneurons that do not degenerate in the *En1*-Het mouse. This differs with mDA neurons that express *En1* and degenerate in the *En1*-Het mouse. EN1 survival activity is therefore independent of its cell autonomous or non-cell autonomous activity, but rather may reflect a higher sensitivity to stress of mDA neurons and αMNs, compared to V1 interneurons and γMNs.

The differential susceptibility of αMNs and γMNs to reduced *En1* expression in the *En1*-Het mouse or to EN1 extracellular availability following scFvEN1 viral expression may be explained by their different functions and the underlying anatomy (Kanning *et al*, 2010). Alpha-MNs not only receive excitatory inputs from local spinal circuits but also receive an important contribution from descending upper motor neurons. They innervate extrafusal fibers and the innervation ratio (i.e., the number of muscle fibers per αMN axon) in humans can reach up to 2000 (Masson *et al*, 2014; Feinstein *et al*, 1954). Gamma-MNs innervate muscle spindles, receive their excitatory inputs from the reticular formation and their inhibitory ones from relay neurons mainly found in the dorsal horn that does not express EN1. They fire after αMNs, at a slower rate and, in humans, the innervation ratio is close to parity (i.e., 1-3 muscle fibers per axon), suggesting that their metabolic demand is less than that of αMNs. Finally, while γMNs can take up exogenous EN1 similarly to αMNs, they do not receive contacts from EN1-expressing interneurons, and thus the functions of EN1 might differ between these two MN types. A full understanding of these distinct functions requires further studies.

The deleterious phenotype is stronger in the *En1*-Het mouse than following extracellular EN1 neutralization in WT mice. This may be due to the fact that EN1 neurotrophic activity exerted on αMNs is purely non-cell autonomous and that scFvEN1 neutralizes less than 50% of extracellular EN1 (due to affinity, levels of expression and/or localization of the antibody). However, the more than twofold reduction in EN1 content in MNs, following scFvEN1 expression (Figure 4), rather suggests a more complex explanation whereby, while scFvEN1 only targets pure non-cell autonomous EN1 activity, the *En1*-Het phenotype may result from a 50% loss in secreted EN1 and, in addition, from cell autonomous dysfunctions affecting V1 interneurons but not inducing their death. V1 interneurons are an essential component of the reciprocal inhibitory αMN circuit and loss of their inhibitory input places αMNs at risk (Quinlan, 2011; Wang *et al*, 2008; Ramírez-Jarquín & Tapia, 2018; Ramírez-Jarquín *et al*, 2014). Furthermore, deficits in inhibitory interneurons in the spinal cord of motoneuron disease models have been reported (Hossaini *et al*, 2011; Allodi *et al*, 2021) and Renshaw cell pathology has been observed following αMN loss (Chang & Martin, 2009; Allodi *et al*, 2021). One possibility is that EN1 regulates the expression of classical trophic factors in spinal V1 interneurons that are secreted and support αMN survival, in conjunction with EN1 signaling (Figure 10). Such synergy between classical secreted factors and HP signaling has been reported for Engrailed and DPP in the fly wing disk (Layalle *et al*, 2011), for EN2, adenosine, and EphrinA5 in chick RGC growth cone navigation (Wizenmann *et al*, 2009; Stettler *et al*, 2012), and for PAX6 and Netrin1 in the migration of chick oligodendrocyte precursors (Di Lullo *et al*, 2011).

**Figure 10.**
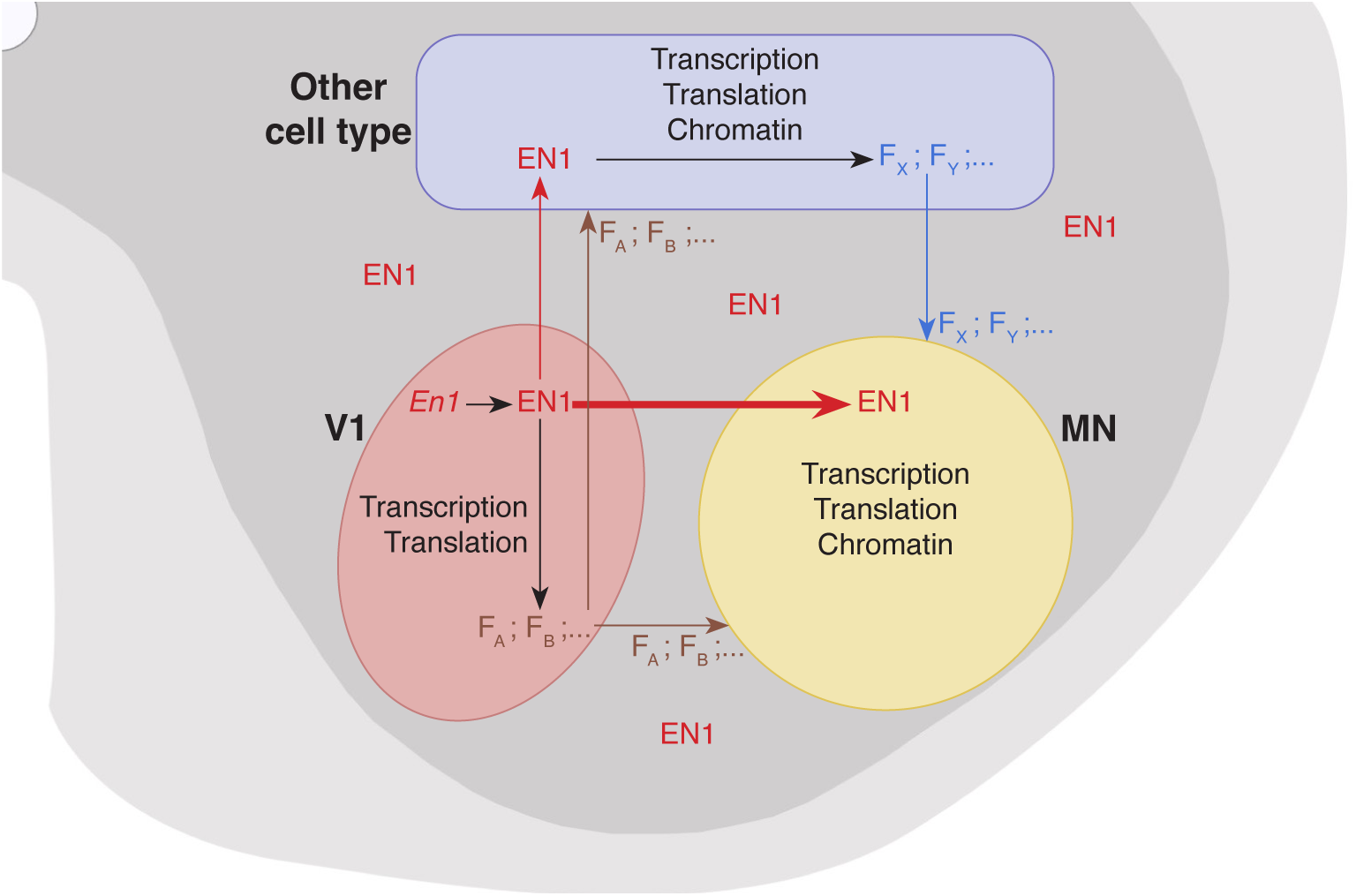
Proposed schema of EN1 cell autonomous and non-cell autonomous activities. EN1 transcribed and synthesized in V1 (pink) has a cell autonomous activity in these cells contributing to the expression of several V1 factors (F_A_, F_B_, …) that may signal to MNs (yellow) and other cell types (blue). All the latter F factors do not necessarily require EN1 activity. EN1 after its secretion by V1 interneurons is preferentially internalized by MNs (fat red arrow) and less by other cell types (thin red arrow), if at all. These other cell types express other F Factors (F_X_, F_Y_, …) which are secreted and are, or not, under non-cell autonomous EN1 activity. As a result, MNs experience the signaling activity of F factors (possibly EN1-dependent as in the scheme, but not necessarily so) and internalize EN1. Following internalization, EN1 can work alone or in synergy with the F factors to regulate transcription, translation and chromatin conformation within MNs.

It remains that the scFvEN1 experiment demonstrates an EN1 non-cell autonomous activity. This activity could be through extracellular EN1 acting directly on αMNs but also indirectly through other cell types, such as astrocytes, oligodendrocytes, or microglial cells (Figure 9). However, the internalization preference of injected EN1 by MNs and the fact that injected EN1 totally restores a healthy phenotype in the *En1*-Het mouse strongly suggest that EN1 exerts its non-cell autonomous activity primarily on αMNs. This activity is reminiscent of the direct non-cell autonomous protective activity of OTX2 for RGCs upon NMDA excitotoxic insult (Torero-Ibad *et al*, 2011). Moreover, when comparing the phenotypes of the *En1*-Het and scFvEN1 mice, one must be aware of the fact that, in the *En1*-Het, EN1 levels are lower very early during development. We thus cannot exclude that extracellular EN1 acts in conjunction with compromised interneuron physiological properties in *En1*-Het animals, even if the interneurons are present in the appropriate number.

EN1 may protect αMNs through one or several of the mechanisms previously characterized for mDA neurons of the SNpc, which can be protected from oxidative stress by a single EN1 injection (Rekaik *et al*, 2015; Blaudin de Thé *et al*, 2018; Thomasson *et al*, 2019). Among these mechanisms are the regulation of Complex I mitochondrial activity, DNA break repair, heterochromatin maintenance, and the repression of long interspersed nuclear elements (LINE-1) expression (Alvarez-Fischer *et al*, 2011; Rekaik *et al*, 2015; Blaudin de Thé *et al*, 2018). The evaluation of these possibilities in the spinal cord will be the objective of future studies but the long-lasting effect of a transient twofold increase in EN1 concentration within MNs strongly suggests an activity taking place at the chromatin level.

The *En1-Het* mouse presents muscle weaknesses, abnormal spinal reflex, NMJ denervation, and αMN loss, all of which are phenotypes reminiscent of changes in ALS patients and in some ALS mouse models. This raises the question as to whether the results reported here are relevant for αMN diseases. To our knowledge, although they may exist in some sporadic cases, as for PD (Haubenberger *et al*, 2011), no studies so far have associated ALS with mutations affecting *En1* integrity or *En1* expression. It is thus clear that the *En1-het* mouse cannot, as yet, be considered as an ALS mouse model. However, *En1* genetic interactions with pathways leading to ALS, and its therapeutic activity in classical mouse and non-human primate models of PD (Alvarez-Fischer *et al*, 2011; Blaudin de Thé *et al*, 2018; Rekaik *et al*, 2015; Thomasson *et al*, 2019), allows us to envisage its use as a long-lasting therapeutic protein in ALS mouse models or iPSC-derived human MNs.

One main mechanism possibly linking EN1 expression to ALS is that, in ALS as in many neurodegenerative diseases, ageing is a major risk factor. Indeed, our results indicate that SQSTM1/p62 expression is a potential marker of age, both in αMNs and γMNs, in agreement with its interaction with redox regulation pathways (Hensley & Harris-White, 2015). Since *En1*-Het mDA neurons are more sensitive than WT to oxidative stress and given that EN1 protects them against experimental oxidative stress (Rekaik *et al*, 2015), we speculate that the up-regulation of SQTSM1/p62 expression, either in the mutant or following extracellular loss due to scFvEN1 expression, reveals an EN1 anti-ageing activity that may explain the ALS-like phenotype of *En1* hypomorphs. This hypothesis is supported by the finding that hEN1 injection at 1 month prevents the upregulation of p62/SQTSM1 in the MNs of 3-month-old En1-Het mice and prevents their loss of strength. Indeed, we now need to follow the effects of gain and loss of EN1 functions in *bona fide* ALS animal models.

In conclusion, we have demonstrated a novel non-cell autonomous activity for EN1 in adult spinal cord. Constitutive reduction in EN1 expression or local neutralization of extracellular EN1 cause motor endplate degeneration, muscle weakness, and αMN loss. This is an example of an important non-cell autonomous activity for this transcription factor as is also the case for EN2, OTX2, PAX6, and VAX1 (Prochiantz & Di Nardo, 2015; Di Nardo *et al*, 2020; Brunet *et al*, 2007; Di Nardo *et al*, 2018). Given that the sequences necessary for intercellular transfer are conserved in most HPs (Prochiantz & Joliot, 2003; Joliot & Prochiantz, 2004) and that such transfer has been demonstrated for about 150 of them (Lee *et al*, 2019), it is not unreasonable to predict that the developmental and physiological non-cell autonomous functions demonstrated for a handful of HPs only represent the tip of the iceberg.

## MATERIALS AND METHODS

### Animal Management

All animal treatments followed the guidelines for the care and use of laboratory animals (US National Institutes of Health), the European Directive number 86/609 (EEC Council for Animal Protection in Experimental Research and Other Scientific Utilization), and French authorizations n° 00703.01 “Therapeutic homeoproteins in Parkinson Disease” and n° APAFIS #6034-2016071110167703 v2, “Spinal cord motoneuron neuroprotection” delivered by the Ministry of higher education, research and innovation. Adult *En1-*Het mice and WT littermates (available Jackson Labs strain #:00712) were housed two to five per cage with *ad libitum* access to food and water and under a 12 h light/dark cycle. Transgenic mouse strain *En1-*Het was bred by the Rodent Breeding Services provided by the Animal Care Services at College de France. Females and males were included in all studies. The endpoint limit for euthanasia was a 15% or greater loss of bodyweight or signs of paralysis; in all experiments, no mouse reached these endpoints.

### Behavior analyses

Mice were habituated to the behavioral room and to the experimenter 24 hours before the day of testing and again before each behavioral test. All tests were performed on the same day and behavioral assessment was carried by evaluators blind to genotype and treatment.

#### Forepaw Grip Strength

The Transducer (IITC Life Science Grip Strength Meter, ALMEMO 2450 AHLBORN, World Precision Instruments) was calibrated and the scale of values set to grams. Each mouse was lifted by the tail to position the front paws at the height of the bar (about 15 cm) and then moved towards the bar. When a symmetric firm grip with both paws was established, the mouse was pulled backward at a constant speed until the grasp was broken and the maximal value was recorded. The test was repeated 5 times per animal with a minimal resting time of 5 minutes between tests and the mean of all values was normalized to the weight of each animal.

#### Inverted Grid Test

The mouse was placed on a wire grid (15×10 cm) and allowed to explore it. After 3-5 minutes, the grid was raised 30 cm above a soft surface and slowly inverted. Latency to release was recorded three times per mouse with a minimum resting time of 5 minutes between trials. The longest latency was used for analysis.

#### Hindlimb Extensor Reflex

Mice were suspended by the tail at a constant height (about 15 cm) and scored for hindlimb extension reflex. The scores were assigned from 0 to 3 as follows: 3 for normal symmetric extension in both hind limbs without visible tremors; 2.5 for normal extension in both hind limbs with tremor in one or both paws; 2.0 for unequal extension of the hind limbs without visible tremors; 1.5 for unequal extension in the hind limbs with tremors in one or both paws, 1.0 for extension reflex in only one hindlimb, 0.5 for minimum extension of both hindlimbs, 0 for absence of any hindlimb extension.

### Tissue preparation

#### Spinal cord

Adult mice were euthanized by a 1 µl/g body weight dose of Dolethal (Euthasol: 4 μg/μl). Spinal cords were dissected and placed in Phosphate Buffer Saline (PBS) to remove meninges and surrounding connective tissues. Cervical and lumbar enlargements were separately placed in paraformaldehyde 4% (PFA, Thermo Scientific) for 1 hour at room temperature (RT) with gentle mixing, washed in PBS three times for 30 minutes at RT and placed in PBS with 20% sucrose overnight at 4°C. After cryoprotection, the tissue was embedded in Tissue Freezing Medium (TFM, Microm Microtech), frozen on dry ice and 30 μm sections were prepared using an HM 560 Microm cryostat (Thermo Scientific).

#### Muscle

The lumbrical muscles were dissected in cold PBS, fixed at RT in 4% PFA for 10 minutes and washed in PBS. Muscle whole-mounts were processed *in toto* to visualize the entire innervation pattern and allow for a detailed NMJ analysis (Sleigh *et al*, 2014).

#### Cresyl violet staining

Slides with 30 μm spinal cord sections were washed in PBS 3 times, cleared in O-Xylene (CARLO-HERBA) for 5 minutes, then hydrated in graded alcohol with increasing water and placed in Cresyl Violet acetate (MERCK). Sections were then dehydrated in increasing alcohol and mounted in Eukitt® Quick-hardening mounting medium (Sigma).

#### Spinal cord and muscle immunofluorescence labeling

Slides with 30 μm spinal cord sections or whole-mount muscles were washed in PBS and permeabilized with 2% Triton. After 30 minutes at RT in 100 μM glycine buffer, 10% Normal Goat Serum (NGS, Invitrogen) or Fetal Bovine Serum (FBS, Gibco) was added in the presence of 1% Triton before incubation with primary antibodies [sheep anti-Choline Acetyltransferase (ChAT) ABCAM 1:1000, goat anti-ChAT Millipore 1:500, rabbit anti-EN1 86/8 1:300 (Sonnier *et al*, 2007), rabbit anti-EN1 LSBio (CliniSciences) 1:200, mouse anti-neurofilament 165kDa Developmental Studies Hybridoma Bank 1:50, mouse anti-synaptic vesicle glycoprotein 2A DSHB 1:100 and rabbit anti-p62 ABCAM 1:1000] overnight at 4°C, washed and further incubated with secondary antibodies for 2 hours at RT. Where indicated, anti-EN1 LSBio activity was neutralized by a 1-hour (RT) preincubation with hEN1 (1.5 EN1/LSBio molar ratio). For muscle staining, α-bungarotoxin (Alexa Fluor 488 conjugate) was included at the same time as the secondary antibodies. Slides were washed and mounted with DAPI Fluoromount-G® (Southern Biotech). Controls without primary antibodies were systematically included.

### RT-qPCR

Spinal cords were removed as above and lumbar enlargements were rapidly frozen on dry ice. Total RNA was extracted (RNeasy Mini kit, Qiagen) and reverse transcribed using the QuantiTect Reverse Transcription kit (Qiagen). RT-qPCR was done using SYBR-Green (Roche Applied Science) and a Light Cycler 480 (Roche Applied Science). Data were analyzed using the “2-ddCt” method and values were normalized to *Glyceraldehyde 3-phosphate dehydrogenase* (*Gapdh*). The following primers were used: *Engrailed-1* sense: CCTGGGTCTACTGCACACG, antisense: CGCTTGTTTTGGAACCAGAT; *Gapdh* sense: TGACGTGCCGCCTGGAGAAAC, antisense: CCGGCATCGAAGGTGGAAGAG.

### Protein and single-chain antibodies

#### Protein

Human EN1 (hEN1) was produced as described (Toreo-Ibad *et al*, 2011) and endotoxins were removed by Triton X-144 phase separation. In brief, pre-condensed 1% Triton X-144 (Sigma) was added to the protein preparation. The solution was incubated for 30 minutes at 4 °C with constant stirring, transferred to 37 °C for 10 minutes and centrifuged at 4000 rpm for 10 minutes at 25 °C. The endotoxin-free protein was aliquoted and kept at -80 °C.

#### Single-chain antibodies

The anti-EN1 single chain antibody plasmid was prepared from the anti-Engrailed 4G11 hybridoma (Developmental Hybridoma Bank, Iowa City, A, USA). Cloning was as described (Wizenmann *et al*, 2009) with or without signal peptide (Lesaffre *et al*, 2007) with 6 myc tags at the C-terminus of the antibody followed by a GFP downstream of a P2A skipping peptide. Addition of a GFAP promoter, insertion in an AAV8 backbone and production of the AAV8 were done by Vector Biolabs.

### Intrathecal injections

Mice were anesthetized with 1 μl/g ketamine-xylazine (Imalgene1000: 10 μg/μl, Rompur 2%: 0.8 μg/μl) and placed on the injection platform. The tail of the animal was taken between two fingers and the spinal column was gently flattened against a padded surface with the other hand. The L3 vertebral spine was identified by palpation and a 23G x 1” needle (0.6×25 mm Terumo tip) was placed at the L1 and T13 groove and inserted through the skin at an angle of 20° (Hylden & Wilcox, 1980). The needle was slowly advanced into the intervertebral space until it reached the injection point, provoking a strong tail-flick reflex. Injections of 5 µl were done at a rate of 1 μl/min with 1 µg recombinant protein, AAV8-IgK-scFvMUT-6xMyc-2A-eGFP (scFvMUT), or AAV8-IgK-scFvEN1-6xMyc-2A-eGFP (scFvEN1) (Vector Biolabs, 10^12 GC/ml). The peptides (ProteoGenix) were co-injected with hEN1 at a 20-fold molar excess of peptides. The needle was left in place for two minutes after injection and then slowly removed. Animals were placed in a warmed chamber until recovery from anesthesia. Extensor reflex and gait analysis were examined 2 and 24 hours after injection to verify the absence of spinal cord damage.

### RNAscope Fluorescent in Situ Hybridization

Mice aged 4.5 months were euthanized with Dolethal and transcardially perfused with PBS followed by 4% PFA in PBS. The lumbar region of the spinal cord was dissected, fixed overnight in 4% PFA at 4 °C and cryoprotected in PBS 30% sucrose for 24 hours at 4 °C. Spinal cord sections 40 µm-thick were prepared using a Leitz (1400) sliding microtome. Sections were washed in PBS, incubated with RNAscope hydrogen peroxide solution from Advanced Cell Diagnostics (ACD) for 10 min at RT, rinsed in Tris-buffered saline with Tween (50 mM Tris-Cl, pH 7.6; 150 mM NaCl, 0.1% Tween 20) at RT, collected on Super Frost plus microscope slides (Thermo Scientific), dried at RT for 1 hour, rapidly immersed in ultrapure water, dried again at RT for 1 hour, heated for 1 hour at 60°C and dried at RT overnight. The next day, sections were immersed in ultrapure water, rapidly dehydrated with 100% ethanol, incubated at 100 °C for 15 minutes in RNAscope 1X Target Retrieval Reagent (ACD), washed in ultrapure water and dehydrated with 100% ethanol for 3 minutes at RT. Protease treatment was carried out using RNAscope Protease Plus solution (ACD) for 30 minutes at 40 °C in a HybEZ oven (ACD). Sections were then washed in PBS before *in situ* hybridization using the RNAscope Multiplex Fluorescent V2 Assay (ACD). Probes were hybridized for 2 hours at 40 °C in HybEZ oven (ACD), followed by incubation with signal amplification reagents according to the manufacturer’s instructions. Probes were purchased from ACD: Mm-*En1*-C1 (catalog #442651), Mm-*Chat*-C2 (catalog #408731-C2), Mm-*Calb1*-C3 (catalog #428431-C3). The hybridized probe signal was visualized and captured on an upright Leica CFS SP5 confocal microscope with a 40x oil objective.

### RNAscope and immunofluorescence co-detection

Sections prepared as described above were collected on Super Frost plus microscope slides and heated for 30 min at 60 °C. RNAscope was performed according to the manufacturer’s instructions (Multiplex Fluorescent V2 Assay and RNA-Protein Co-detection Ancillary Kit Cat No. 323180). In brief, sections were post-fixed in 4% PFA for 15 min at 4 °C, dehydrated in 50%, 70%, 100% (twice) ethanol solutions for 5 min each, incubated with RNAscope hydrogen peroxide solution for 10 min at RT, rinsed in ultrapure water, incubated at 100 °C for 5 minutes in RNAscope 1X Co-detection Target Retrieval, rinsed in ultrapure water and washed in Phosphate buffer, 0.1% tween 20 pH7.2 (PBS-T). Sections were incubated overnight at 4 °C with anti-EN1 LSBio diluted (1:200) in Co-detection Antibody Diluent, washed in PBS-T, fixed in 4% PFA for 30 min at RT and washed twice in PBS-T. Sections were treated with protease 30 min at 40 °C in a HybEZ oven, washed in PBS before *in situ* hybridization using the RNAscope Multiplex Fluorescent V2 Assay. Probes were hybridized for 2 hours at 40 °C in HybEZ oven, followed by incubation with signal amplification reagents according to manufacturer instructions. Finally, sections were incubated with the secondary antibody diluted in the Co-detection Antibody Diluent for 30 min at RT.

### Western blots

#### Tissue extracts

Spinal cords and ventral midbrains dissected from adult mice were kept frozen at -80 °C. After thawing on ice and 3 rinses in PBS, tissues were resuspended in 500 µl dissociation buffer (DB: 50 mM Tris-HCl pH8.0, 0.3 M NaCl, Triton X-100 1%, EDTA-free Complete Proteases inhibitor cocktail from Roche and 250U/ml of benzonase endonuclease from Sigma). After homogenization in a Dounce tissue homogenizer, 10 times with the A piston and 10 times with the B piston, lysates were incubated for 30 min on ice, sonicated for 5 min in Diagenode Bioruptor set with 30/30 seconds ON/OFF at high power, incubated on ice for 30 min and centrifuged for 10 min at 12000 *g*. Supernatants were harvested, their protein contents measured (Pierce BCA assay) and frozen at -80 °C.

#### Western blots

Invitrogen Nu-PAGE 4-12% acrylamide 1 mm SDS-PAGE gels were used to run spinal cord extracts or the hEN1recombinant protein. Extracts (2-3 µg total proteins) and 2 ng of purified EN1 were diluted in 10 µl Invitrogen LDS-DTT Sample Buffer, heated for 10 min at 70°C, briefly centrifuged at 10000 *g* and loaded. For the immunoprecipitation followed by Western blot, 1/200 of total inputs or Flow-Throughs and 1/20 of the total protein A– sepharose E2 elution were used per well. SeeBlue 2 and molecular weight markers were from Invitrogen. Gels were run in MOPS buffer at 150 V for 1 hour and 15 min and the proteins transferred for 30 min at 100 V on Millipore PVDF membranes (BIORAD Criterion Blotter apparatus). Membranes were blocked for 1 hour at RT with 4% non-fat dry milk (NFDM) diluted in TBS containing 0.2% Tween-20 (TBST). Antibodies to EN1 86/8 and LSBio were diluted 1/200 in TBST 4% NFDM and incubated overnight at 4 °C. Blots were rinsed 3 times for 15 min each with TBST and incubated with HRP-Labeled anti-Rabbit antibody diluted 1/2500 for 1 hour at RT. When preceded by immunoprecipitations, revelation was with HRP-protein G diluted 1/5000 for 1hour at RT followed by 3 TBST rinses of 15 min each. HRP activity was revealed with chemiluminescence substrate CLARITY from BIORAD and the light was quantified with the FUJI LAS 4000 imager. Signals were quantified with the Image Studio Lite software from Li-Cor.

### Bioinformatic analysis

Differentially expressed genes (cutoff p<0.05) from the RNA-seq performed on microdissected six-week-old SNpc from WT and *En1*-*Het* mice (GSE72321) (Rekaik *et al*, 2015) were compared to genes identified from an RNA-seq of microdissected MNs from WT mice (GSE38820)^41^. Transcripts present in the two lists were retained. Using STRING database, 10 biological processes of interest were selected including regulation of gene expression, cellular response to stress, cell communication, regulation of I-kappaB kinase/NF-kappaB signaling, cellular response to DNA damage stimulus, response to oxygen-containing compounds, locomotion, DNA repair, modulation of chemical synaptic transmission, and brain development. Using these pathways reduced the list of selected genes to 402. To the 402 genes, we used the STRING database to identify known interactions with the four main genes involved in familial ALS (fALS): *SOD1*, *FUS, C9Orf72* and *TARDBP*.

### Image analyses

Cresyl violet-stained spinal cord section images were acquired with a Nikon-i90 microscope under bright field conditions at 10x with a Z-stack step of 0.5 μm. RNAscope FISH images were acquired using a Yokogawa X1 Spinning Disk confocal microscope at 20x (Nikon) and acquisitions of 3D z-stack (1 μm) using 491 nm, 561 nm and 642 nm laser beams. For cell quantification, at least five lumbar spinal cord sections (10 ventral horns) through levels L1 to L5 were analyzed for each animal. Immunofluorescence-stained spinal cord sections images were acquired with a Leica SP5 inverted confocal microscope at 20x (Leica DMI6000) and acquisitions of 3D z-stack (0.5 μm) were made using the UV (405 nm, 50 mW), Argon (488 nm, 200 mW) and DPSS (561 nm, 15 mW) lasers. For cell counting, at least five 30 μm thick spinal cord sections separated by ≥ 900μm were analyzed for each animal. ChAT+ cells and Cresyl violet-stained cells with a soma area above 100μm^2^ were manually outlined using ImageJ software (NIH, Bethesda, MD) and their area determined. Analyses were carried out on Z-stacks through the entire 30μm thickness of the section. Cells were classified as small (100-199 μm^2^ cross sectional area), intermediate (200-299 μm^2^) or large (> 300 μm^2^). For example, in a WT mouse 4.5 months of age 558, 158 and 112 cells were counted in the 100-199, 200-299 and >300 µm2 classes, respectively (5 sections). In a 4.5-month-old *En1*-Het mouse, the values were 562, 149, and 66, respectively. The data in the graphs are the average number of each cell category in one ventral horn. Thus, the WT mouse had an average of 11.2 large MNs and the En1-*Het* mouse had an average of 6.6 large MNs. ChAT and Cresyl violet gave similar results for medium size (200-299 µm^2^) and large (>300 µm^2^) neurons thus allowing us to characterize αMNs based on size (>300 µm^2^), with or without additional ChAT staining. For p62 staining analysis, the motoneuron region of five lumbar sections were acquired at 40x for each animal. Mean intensity was measured in ChAT+ cells with a soma greater than 150 μm^2^ using ImageJ software. For each experiment, image acquisition was performed with the same parameter settings of the confocal microscope to allow for comparison between experimental conditions.

For endplate analysis, lumbrical muscles were imaged with a Leica SP5 inverted confocal microscope (Leica DMI6000) with a motorized XY stage. 3D z-stack (0.5 μm) acquisitions were made as above (UV, Argon and DPSS) and images analyzed using ImageJ software. Analyses were performed blind to the genotype and treatment. The Z-stack with maximum overall intensity was determined. Red (SV2A/NF) and green channels (α-BGT) were separated and images were filtered for better segmentation using a Median Filter with a radius of 2 pixels. Each channel image was then converted to a gray LUT and an automatic threshold was established for both channels to outline the areas of analysis. For postsynaptic analysis, each endplate was identified by α-BGT signal. The integrated density of one channel overlapping the other channel was determined using the AND function in the ROI manager for each pair of measurements. Measurements were performed in the original images and the overlap was quantified based on integrated density as a percentage (overlap values/ total values in the α-BGT channel x100). Endplates were categorized as fully innervated, partially innervated, or denervated. A fully innervated endplate is defined as an endplate in which 80% or more of the green pixels (α-BGT) are covered by a red pixel (SV2A/NF), a partially innervated one is between 20 and 80% and a denervated one below 20% coverage. Endplate morphology was evaluated by counting the number of endplates with perforations (areas lacking α-bungarotoxin staining). All analyses were performed on the entire Z-stacks through the NMJ.

### Statistical analyses

Sample size estimate was based on our previous observations^15^. Results are expressed as mean ± SD. Statistical significance was determined as follows. For RT-qPCR, WT and *En1-*Het mice were compared by Unpaired T-test with equal SD. For behavioral and NMJ analyses and αMN counting in the time-course studies a 2-way ANOVA was performed and if the main effect was significant groups were compared by Unpaired T-test with equal SD comparing WT with *En1-*Het for each time point. For intrathecal injections, behavioral and NMJ analyses and αMN counting, experimental data were compared by one-way ANOVA followed by a post hoc Tukey test for comparisons to WT. For behavioral analysis in the time-course protection of injected hEN1 a 2-way ANOVA was performed and if the main effect was significant groups were compared by Unpaired T-test with equal variances comparing WT with *En1-*Het injected at each time point.

The p62 measures are bounded between 0 and an upper quantification limit (ceiling effect), and classical normally distributed models are ill-suited to account for these two constraints. We solicited PharmaLex Belgium to develop a suitable analysis. Since it can take a wide variety of shapes and is naturally bounded in the [0;1] interval, beta-distribution was fitted on the p62 measure, for all MNs, γMNs, and αMNs separately, after rescaling the original data to the [0;1] interval. The shape and scale parameters account for time, group, and time*group interaction as fixed effect. It models the variance observed in the data by time, group, and time*group interaction, meaning each combination has its own variability. Finally, the mouse-to-mouse variability (or between mouse variability) is modeled by time and group.

## FIGURE LEGENDS

## Acknowledgements

We would like to thank Anne Bousseau, Ariel Di Nardo, Lizzie di Lullo and Julien Spatazza for reading the manuscript and discussing experiments and results with the authors. We are grateful for the help brought by the CIRB Imaging and animal facilities. Funding was provided by Memolife Labex PhD fellowship for SEVA, Association Nationale de la Recherche et de la Technologie (Cifre/ANRT n°2017/0488) to ML, BrainEver, Homeosign ERC-AdG n°339379 and Fondation Schueller-Bettencourt.

Author contributions: ML and SEVA contributed to experiment design and execution; E.P-H developed the single-chain antibodies; L.J-A participated in all histochemistry experiments; ED did the immunoprecipitation and western blot experiments; KLM and AP designed the research and wrote the manuscript.

Conflict of interest statement: AP and KLM are co-founders and hold shares in BrainEver, a company developing HPs for therapeutic use.

Funding sources: Memolife Labex PhD fellowship for SEVA, Association Nationale de la Recherche et de la Technologie (Cifre/ANRT n°2017/0488) to ML, BrainEver, Homeosign ERC-AdG n°339379, Fondation Schueller-Bettencourt.

---

**Figure EV1.**
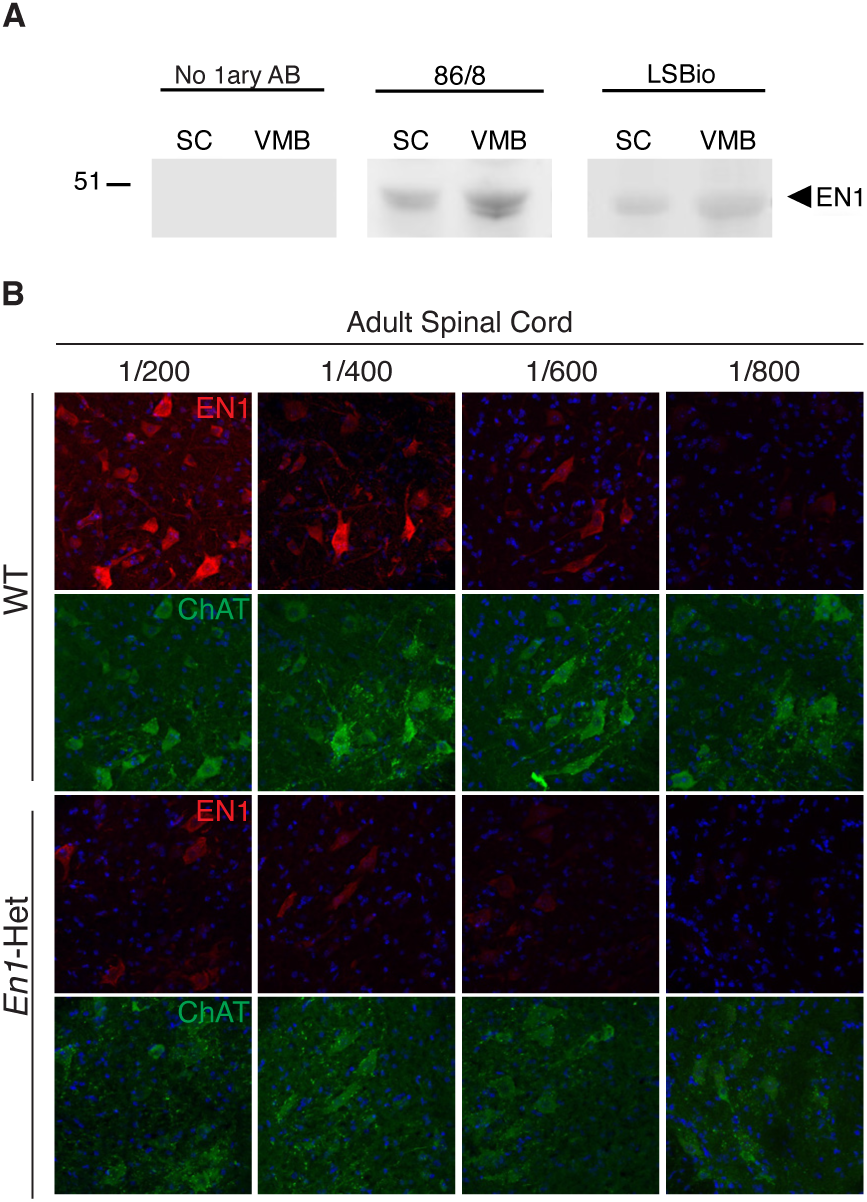
Characterization of the anti-EN1 LSBio antibody. **A.** Western blots of spinal cord (SC) and ventral midbrain (VMB) extracts demonstrating that the 86/8 and LSBio antibodies recognize in both structures the same protein migrating with recombinant EN1 velocity. No staining is observed in absence of primary antibody (left panel). This experiment was performed twice. **B.** Double-staining of 3-month-old WT and *En1*-Het ventral MNs with the anti-ChAT antibody and the anti-EN1 LSBio antibody at various dilutions. EN1 staining decreases with increasing dilutions of the antibody. The loss of staining is more rapid in *En1*-Het than in WT mice. This experiment was performed once.

**Figure EV2.**
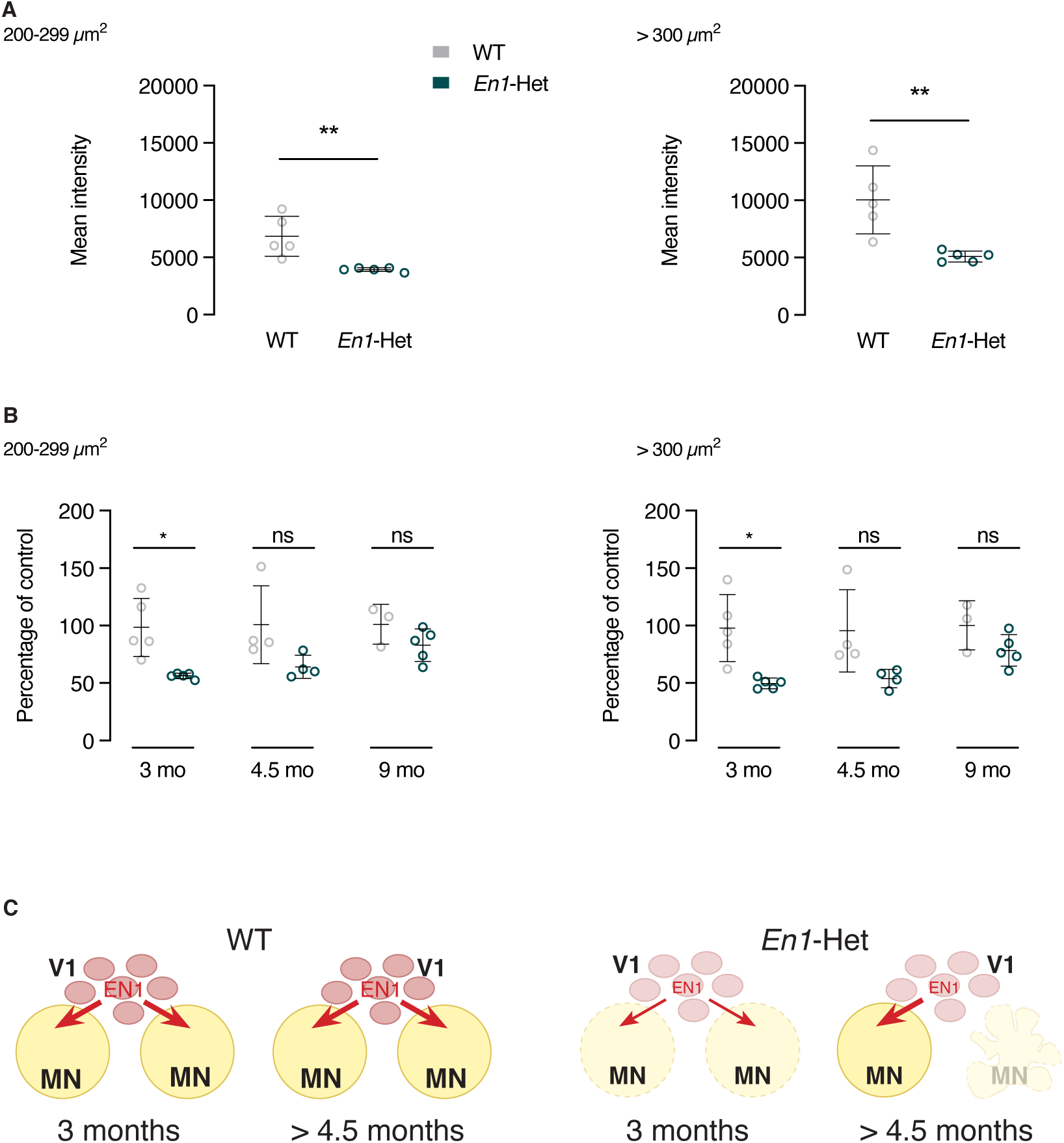
Evolution of EN1 content per MN in WT and *En1*-Het mice. **A.** Analysis of EN1 amount in MNs at 3 months in WT and *En1*-Het mice. EN1 content is reduced by about half in γMNs (left panel) and αMNs (right panel). EN1 was revealed by the LSBio antibody allowing for the visualization of endogenous EN1. Unpaired two-sided t-test with equal SD (**p<0.005; n=5). **B.** Quantification of EN1 amount in γMNs and αMNs with the LSBio antibody at 3, 4.5 and 9 months in WT and *En1*-Het mice. Values at 3 months correspond to the ones showed in panel A. At 4.5 and 9 months, the amount of EN1 in MNs is similar in *En1*-Het and WT mice suggesting that, with time, each remaining MN receives a higher amount of EN1. Two-way ANOVA showed a significant main effect for the γMNs (F(1,20)=16.93 p=0,0005) and αMNs (F(1,20)=19.03, p=0.0003). Unpaired two-sided t-test with equal SD (*p<0.05; n=4-5). This experiment was performed once. **A. C.** Hypothetical representation of EN1 availability to αMNs in WT and Het mice. In the *En1*-Het mouse, each V1 interneurons only provides half as much EN1 to the full population of αMNs at 3 months of age. At 4.5 months of age and later, half of the αMNs have been lost allowing each remaining αMN to receive its full complement of EN1 from the V1 interneurons.

**Figure EV3.**
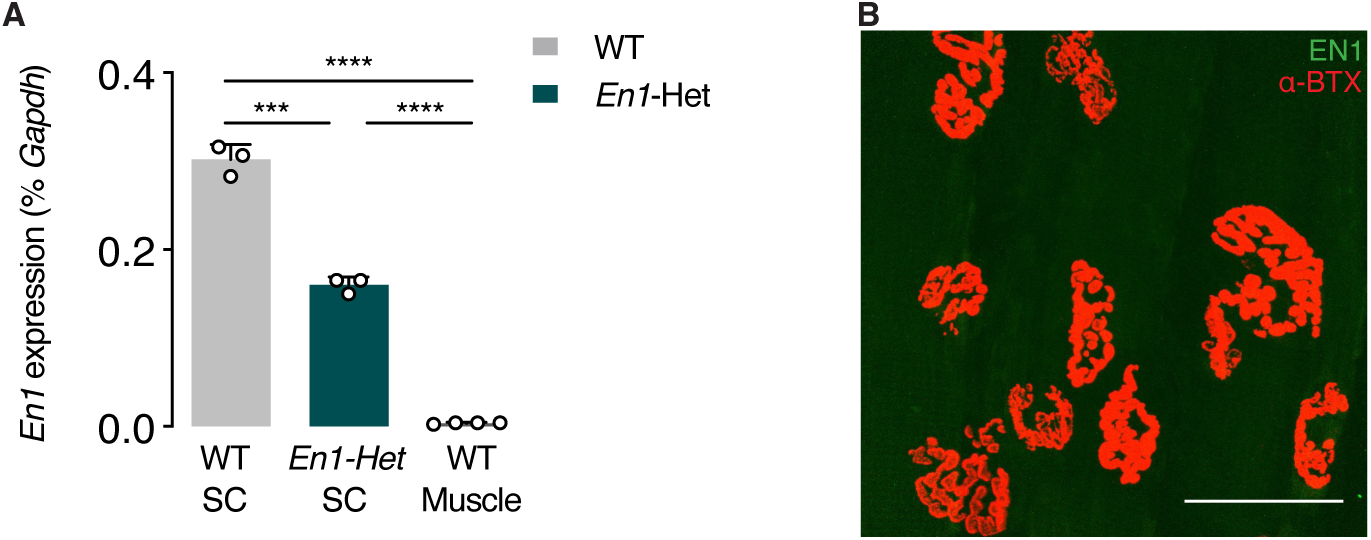
Absence of *En1* transcription or EN1 protein at endplate level. **A.** RT-qPCR of RNA from the lumbar enlargement at 4.5 months of WT and *En1*-Het mice and from WT muscle. *En1* expression is absent from the muscle of WT mice. Unpaired two-sided t-test with equal SD (***p<0.005; ****p<0.0005; n=4-5). **B.** Immunohistochemistry for EN1 protein (LSBio antibody, in green) shows its absence at the level of the NMJ (α-BTX, in red). This experiment was done once.

**Figure EV4.**
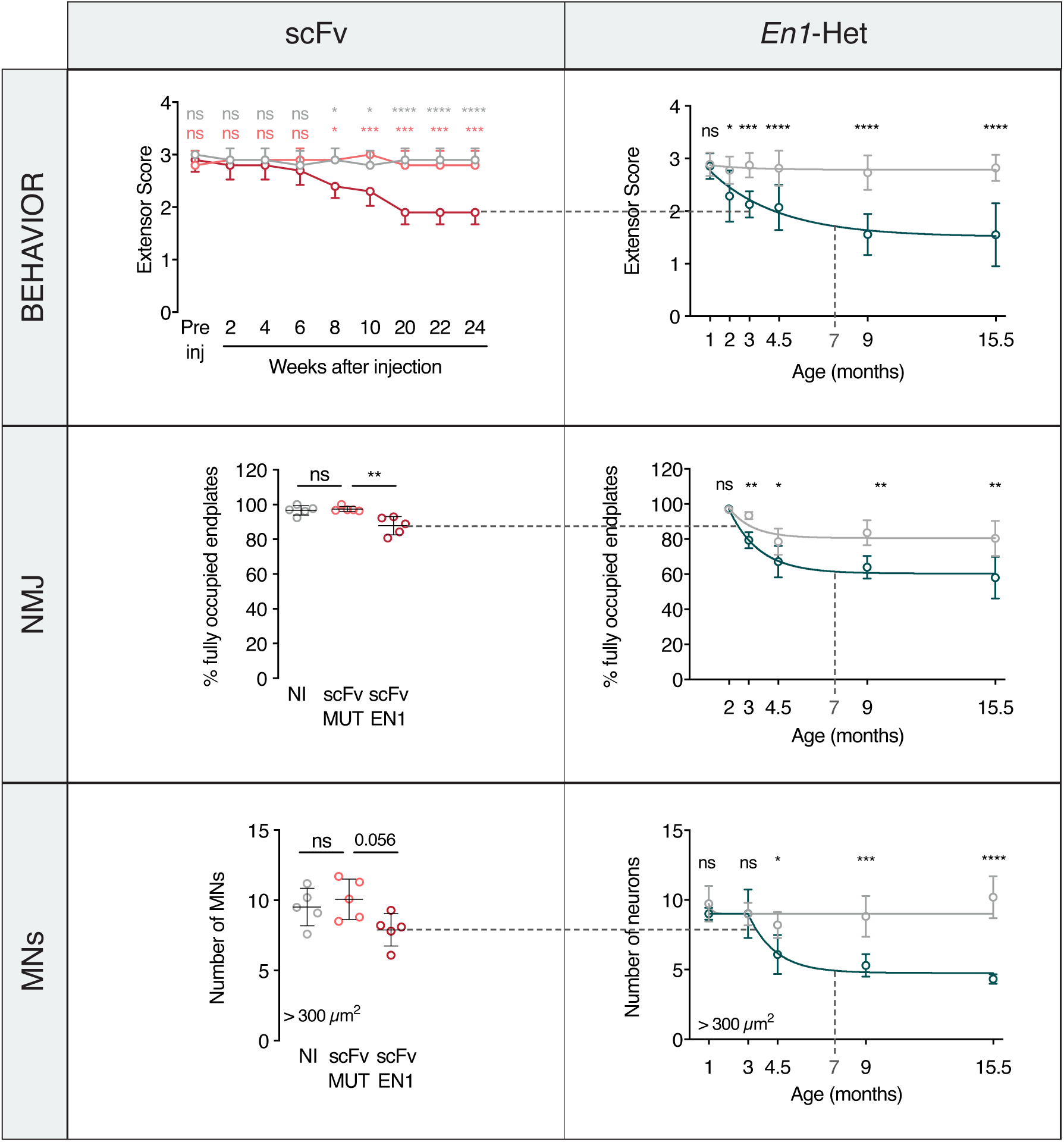
Comparison of *En1*-Het and scFvEN1-expressing mice phenotypes. Comparison between *En1*-het mouse and scFvEN1 models. Data and graphs are from main figures (primarily Figures 2 and 4). WT mice injected with scFvEN1 show similar result to those obtained in the *En1*-het mouse with a milder strength loss, a smaller decrease in the number of fully-occupied endplates, and the specific loss of large size αMNs. At 7 months, scFvEN1 injected mice have a phenotype similar to 3-month-old *En1*-het mice.

